# Human deep sleep facilitates faster cerebrospinal fluid dynamics linked to brain oscillations for sleep homeostasis and memory

**DOI:** 10.1101/2024.08.30.610454

**Authors:** Makoto Uji, Xuemei Li, An Saotome, Ryosuke Katsumata, R. Allen Waggoner, Chisato Suzuki, Kenichi Ueno, Sayaka Aritake, Masako Tamaki

## Abstract

How sleep maintains our healthy brain function has remained one of the biggest mysteries in neuroscience, medical settings, and daily lives. While cerebrospinal fluid (CSF) during sleep have been implicated in metabolic waste reduction in animals, how CSF dynamics are driven in the healthy human brain during deep sleep remains elusive. A myriad of research has shown that crucial cognitive processing manifests in slow wave and rapid-eye movement (REM) sleep, suggesting that a key to maintaining brain functions lies in deep sleep. By leveraging a simultaneous sparse-fMRI and polysomnography method, we demonstrate that deep sleep-specific faster CSF dynamics are associated with spontaneous brain oscillations in healthy young human participants. Slow waves and sleep spindles during slow-wave sleep and rapid eye movements and sawtooth waves during rapid eye movement (REM) sleep are tightly linked to low-amplitude faster CSF fluctuations. In contrast, slow waves during light sleep and arousals produced large but slower CSF signal changes. Furthermore, CSF signals are significantly faster in frequency during deep than light sleep. These brain oscillations during light and deep sleep recruited essentially different brain networks, with deep sleep involving memory and homeostatic circuits. Thus, human deep sleep has a unique way of enabling faster CSF dynamics that are distinctive from arousal mechanisms.

**Significance Statement:** Sleep is indispensable to our life, but its functions remain a significant mystery in the field of neuroscience. One of the most enigmatic issues in sleep is whether and how sleep regulates the CSF. The present study demonstrates deep sleep-specific faster CSF dynamics time locked to sleep brain oscillations in healthy young human participants. Slow waves and sleep spindles during slow-wave sleep and rapid eye movements and sawtooth waves during REM sleep are tightly linked to CSF fluctuations, contributing to faster CSF signals. Our results consistently demonstrate that human deep sleep has a unique way of enabling faster CSF dynamics that are distinctive from arousal mechanisms.

## Introduction

Sleep is crucial for the maintenance of healthy brain functions (1–3). Poor sleep results in a wide variety of health problems, including neurodegenerative diseases such as Alzheimer’s disease (3, 4). However, how sleep maintains human brain functions has remained one of the biggest mysteries in neuroscience, medicine, and daily life. Cerebrospinal fluid (CSF) during sleep has recently been proposed as one of the essential mechanisms for supporting brain functions (5–9). Research on animal models has suggested that brain metabolite waste is reduced specifically during sleep (5). This process is proposed to rely on CSF flows to reduce metabolic waste from the brain (6). In rodents, the amount of CSF tracer influx increases during deep nonrapid-eye-movement (NREM) sleep or slow-wave sleep compared to during wakefulness (5). Clearance and CSF movement were facilitated by vasomotion associated with norepinephrine level during sleep (9). Additionally, accumulating body of evidence suggests that CSF during sleep plays an important role in metabolic clearance also in humans. Tracer substances administered to CSF were reduced from the brain after sleep, whereas sleep deprivation impaired the clearance (8). Another study has reported that impaired sleep quality was associated with increased CSF tracer enrichment (7).

While recent findings suggest the contribution of sleep to CSF clearance, how CSF dynamics are maintained in healthy human sleep has yet to be clarified. Due to safety concerns when MRI methods are simultaneously measured with polysomnography (PSG), which is necessary for sleep method that allows measuring CSF dynamics or strengths concurrently with PSG (11–13), while several other MRI techniques have been proposed for measuring CSF dynamics independent of PSG (14, 15). Hereafter we refer to the fMRI signals measured from ventricles as CSF signals. By measuring fMRI simultaneously with PSG during sleep, the slow frequency component (∼0.05 Hz) in the CSF signals was found to be stronger during light sleep than during wakefulness in correlation with EEG delta activity (11). What during light sleep has caused such slow fluctuations has been unclear. Since K-complexes, a large abrupt biphasic EEG activity, and EEG arousals occur frequently during light nonrapid-eye-movement (NREM) sleep (16, 17), neural or physiological activities induced by these events may contribute to the slow CSF signal changes (9, 18). However, crucially, spontaneous brain oscillations and events, including slow waves and sleep spindles, continuously occur during deep sleep (19). If neural activity caused by spontaneous brain activities contributes to modulation of CSF signals, the CSF signals should oscillate in link with these oscillations. Additionally, the CSF signals should oscillate faster during slow-wave sleep than those during wakefulness or light sleep due to the abundancy of these sleep events, especially slow waves, when arousal is largely suppressed (20). These rationales led to our hypothesis that human deep sleep plays a specific role in accelerating CSF signal changes, which should be tightly linked to sleep brain oscillations and events.

We demonstrate that sleep brain oscillations during slow-wave sleep and REM sleep are indeed associated with significant changes in the CSF signals in healthy young human participants. We found that CSF signals fluctuate significantly faster during slow-wave sleep than during light NREM sleep, time-locked to slow waves and sleep spindles. During REM sleep, rapid eye movements and sawtooth waves were associated with the significant CSF signal increase. Since we did not find correlation between the CSF signal changes and any of respiration, heat rates, and head motions, none of respirations, heart rates, or head motions were likely to cause the differences in the CSF signal changes that were found among sleep stages and brain oscillations. We further confirmed that characteristics of brain networks activated during slow-wave and REM sleep are distinct from those during light NREM sleep or arousals. The present study suggests a fundamental property of deep sleep for the maintenance of brain functions and offers a clue as to why deep sleep matters for the human brain.

## Results

One of the challenges in using fMRI in sleep research is the acoustic noise in MRI scans that cause frequent arousals and prolonged wakefulness, inhibiting participants from reaching deeper sleep (10, 21). To acquire slow-wave sleep and REM sleep data efficiently, we utilized a sparse fMRI sequence which had a 2.1 s quiet period between 0.9 s of MRI image acquisitions measured simultaneously with PSG. This protocol was inspired by previous studies that successfully acquired slow-wave sleep and REM sleep data using magnetic resonance spectroscopy (MRS), where the duration of MRI signal acquisition was approximately 1 s, separated by quiet periods (22–24). Leveraging the sparse method (**Fig. S1**; see ***Experimental design*** and ***MRI acquisition*** in **Materials and Methods**), we acquired a total of 50 datasets from twenty-five young healthy participants (mean 23.6 ± 4.8 years old, 15 female) during mid-afternoon nap. The average sleep duration was 69.6 ± 2.9 min (*mean* ± *SE*), including sessions with slow-wave sleep and/or REM sleep (**Table S1**). The fMRI data were denoised using cardiac and respiration signals (25, 26). The CSF signals were extracted from each participant’s lateral ventricles to measure the signal changes caused by CSF, because these are the closest ventricles from the choroid plexi that have been generally regarded as one of the main sources of CSF production (27, 28) (**Fig. 1a**). Any epochs containing arousals (29) or motion artifacts, defined as an epoch containing artifacts that lasted longer than 1s or a participant showing motion in the video recording (see ***Sleep stage scoring*** in **Materials and Methods**), were removed from the analyses (except for the analyses targeting arousals; **Fig. 3a**).

**Figure 1.**
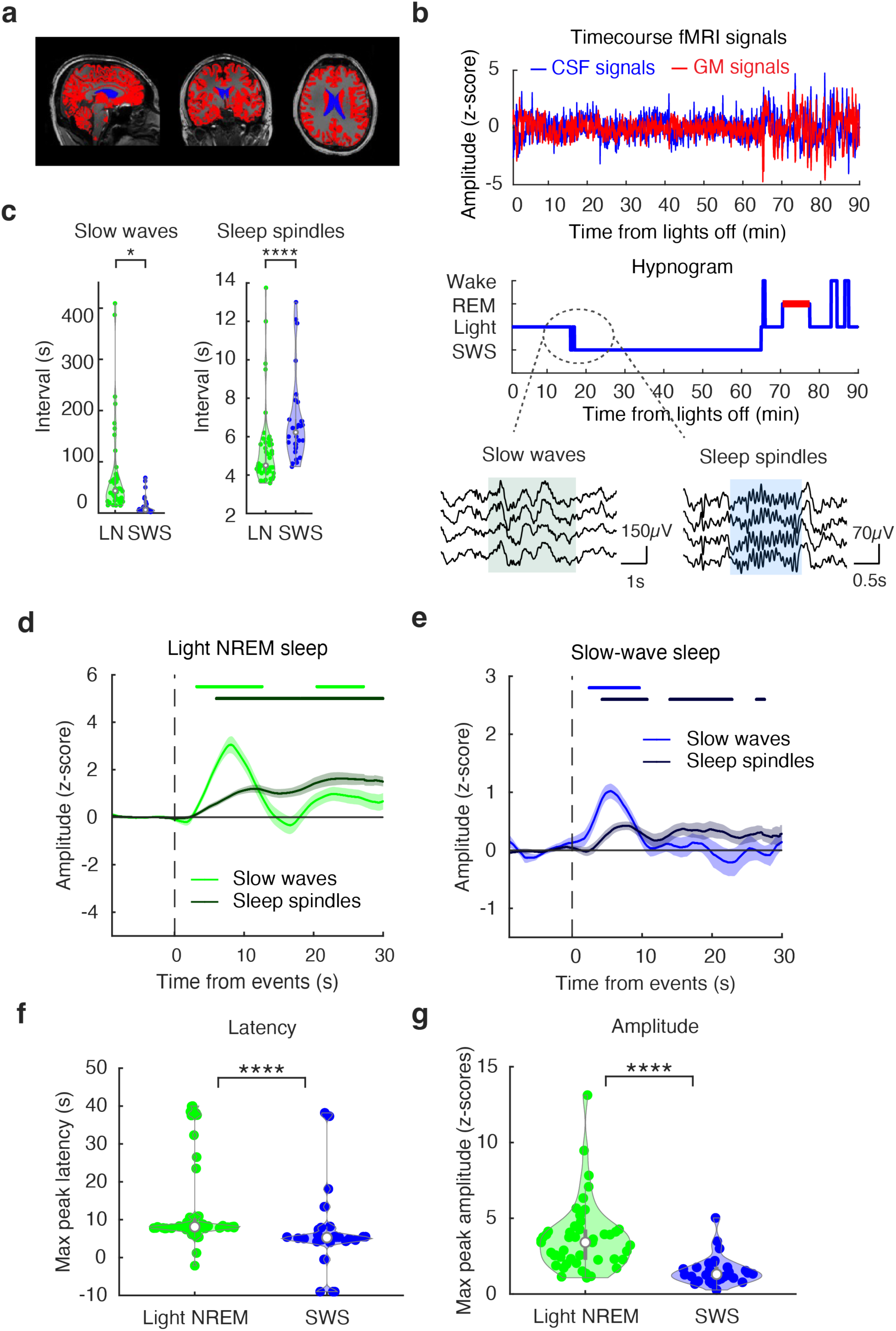
Sleep-specific brain oscillations are followed by faster and milder CSF signal changes during slow-wave sleep. **a**, A schematic of the regions of interest. Blue, lateral ventricles. Red, whole gray matter (GM). **b,** Example MRI signals (top; blue, CSF signals. red, GM signals), corresponding sleep stages (middle) during a 90-min sleep session, and example slow waves and sleep spindles (bottom). Wake, wakefulness. SWS, slow-wave sleep. Light, light NREM sleep. REM, REM sleep. **c,** Intervals of slow waves (left) and sleep spindles (right) during light NREM (green) and slow-wave sleep (blue). Two-tailed Wilcoxson signed-rank tests, \**P* < 0.05, \*\*\*\**P* < 0.001. LN, light NREM sleep. SWS, slow-wave sleep. **d,** CSF signal changes during light NREM sleep time-locked to the onset of slow waves (light green) and sleep spindles (dark green). The horizontal bars (light green, slow waves; dark green, spindles) indicate statistical significance compared with zero baselines via two-tailed one-sample *t*-tests (FDR corrected, *Ps* <0.05). **e,** CSF signal changes during slow-wave sleep time-locked to the onset of slow waves (light blue) and sleep spindles (dark blue). The horizontal bars (light blue: slow waves; dark blue: spindles) indicate statistical significance compared with zero baselines via two-tailed one-sample *t*-tests (FDR corrected *Ps* < 0.05). **f,** Peak latencies of CSF signals during light NREM (green) and slow-wave sleep (blue) induced by slow waves. Two-tailed Wilcoxson signed-rank tests, \*\*\*\**P* < 0.001. **g,** Peak amplitudes of CSF signal changes during light NREM (green) and slow-wave sleep (blue) induced by slow waves. Two-tailed Wilcoxson signed-rank tests, \*\*\*\**P* < 0.001. SWS, slow-wave sleep. See **Supplementary Fig. S2** for the relationship between the CSF and GM signals. SWS, slow-wave sleep.

### Slow waves and sleep spindles accompany CSF signal fluctuations during slow-wave sleep

If sleep brain oscillations are linked to the CSF signal changes, CSF signal peaks should be present when the signals are time-locked to brain oscillations. Since slow-wave sleep is characterized by ample slow waves in addition to sleep spindles (19, 29), the peak latency should be shorter during slow-wave sleep than during light NREM sleep due to the differences in abundancy of these events. To test whether CSF signal changes follow sleep brain oscillations and whether the latency differs between different sleep stages, EEG signals recorded simultaneously with fMRI data had artifacts removed, sleep-stage scored in accordance with American Academy of Sleep Medicine criteria (29). Then slow waves and sleep spindles were detected. Specifically, high-amplitude slow waves with peak-to-peak amplitude of larger than 140 μV (30, 31) and sleep spindles of frequency between 12-15Hz with the duration of 0.5–3s (32) were detected from the EEG signals during light NREM (N2 sleep stage) and slow-wave sleep (N3 sleep stage; for more details see ***Sleep event detection*** and ***fMRI time course analysis*** in **Materials and Methods**, **Fig. 1b**). The CSF signal changes were measured for each event. If sleep brain oscillations are associated with CSF signals, significant changes in the CSF signals would be observed after the onset of oscillations.

The number of slow waves were significantly larger during slow-wave sleep than during light NREM sleep (**Fig. 1c**; mean intervals between events, light NREM sleep, 73.9 ± 13.05 s; slow-wave sleep, 14.8 ± 3.52 s, *mean* ± *SE*; Wilcoxson signed-rank test, two-sided with Bonferroni correction: *n* = 27, *z* = 3.9401, *P* = 8.1448e-05), whereas spindles were detected more frequently during light NREM sleep than during slow-wave sleep (**Fig. 1c**; mean intervals between events, light NREM sleep, 5.3 ± 0.29 s; slow-wave sleep, 6.8 ± 0.41 s, Wilcoxson signed-rank test, two-sided with Bonferroni correction: *n* = 30, *z* = 3.9289, *P* = 8.5348e-05).

CSF signal changes that occurred after brain oscillations exhibit different characteristics depending on the depth of sleep. During light NREM sleep, in addition to sleep spindles that were correlated with significant CSF signal changes at 6 s after onset (**Fig. 1d**, dark green), slow waves preceded a large CSF signal peak at 8 s after onset (**Fig. 1d**, light green). During slow-wave sleep, slow waves triggered CSF signal changes peaking at 5.5 s **(Fig. 1e**, light blue), which was shorter in latency (**Fig. 1f**, Wilcoxson signed-rank test, two-sided with Bonferroni correction: *n* = 28, *z* = 3.0060, *P* = 0.0026) and smaller in amplitude (**Fig. 1g**, Wilcoxson signed-rank test, two-sided with Bonferroni correction: *n* = 28, *z* = 3.8028, *P* = 1.4305e-04) than light NREM sleep. During slow-wave sleep, sleep spindles again preceded CSF signal changes at 4 s after their onset, which lasted for approximately 10 s (**Fig. 1e**, dark blue). The CSF signal changes were significantly correlated with GM signal changes with GM preceding CSF signal changes (**Fig. S2**). These findings demonstrate that CSF signal changes are indeed correlated with brain oscillations during sleep and that deep sleep was characterized by shorter latency and smaller amplitude CSF signal changes that follow GM signal changes.

To further confirm the relationship between the CSF signal changes and sleep brain oscillations, we conducted an inverse analysis; we investigated the EEG topographical changes based on CSF signal positive peak events (see ***CSF peak locked EEG topography analysis*** in **Materials and Methods**). The segmented EEG data were averaged for each sleep stage (**Fig. S3**). The EEG topographic map indicated a negative peak centered around the frontal region, resembling a topography of K-complexes (33, 34), which peaked 8 s *before* the CSF positive peak events during light NREM sleep (note that slow waves during light NREM sleep were followed by a CSF peak 8s *after* the onset of slow wave events, **Fig. 1d**). During slow-wave sleep, a negative peak spread through the midline of frontal to parietal regions resembling those found typically in slow waves (35) which peaked around 5 s *before* the CSF positive peak events during slow-wave sleep (note that slow waves during slow-wave sleep were followed by a CSF peak 5.5s *after* the onset of slow wave events, **Fig. 1e**). These inverse analyses further confirmed that sleep brain oscillations, mainly slow waves, and CSF signal fluctuations are linked.

### CSF signal changes follow rapid eye movements and sawtooth waves during REM sleep

Rapid eye movements and sawtooth waves during REM sleep have been associated with memory processing and conscious experiences (36, 37). Thus, if brain activities during REM sleep, not limited to NREM sleep, contribute to CSF signal changes, CSF signals should show significant changes that are time-locked to these events. To this end, we detected typical REM sleep events, such as rapid eye movements and sawtooth waves observed during REM sleep (***Sleep event detection*** in **Materials and Methods**; **Fig. 2a**). We combined all of the REM sleep events together since the number of events was limited. We found that REM sleep events accompanied the CSF signal fluctuations 30 s after the onset of the events (**Fig. 2b**; see **Fig. S4a & b** for the results for each event). In contrast to NREM sleep events, REM sleep events took a longer time to induce a smaller CSF signal fluctuation (**Fig. 2c**). We next tested whether global brain activation occurs prior to the CSF peaks, similarly to NREM sleep. We found that REM sleep events were also followed by a GM signal increase at 20 s after onset (**Fig. 2d, Fig. S4c & d**), approximately 10 s before the CSF peak. However, REM sleep events showed a significant positive, not negative, correlation between the CSF and GM signals (**Fig. 2e, *n*** = 14, Pearson’s *r* = 0.31 at lag –6.1 s, *P* < 1e-10, *CI* = [0.16, 0.46]), suggesting different mechanisms for CSF and GM coupling between NREM and REM sleep (see also **Fig. S2c & d** for CSF and GM coupling during NREM sleep).

**Figure 2.**
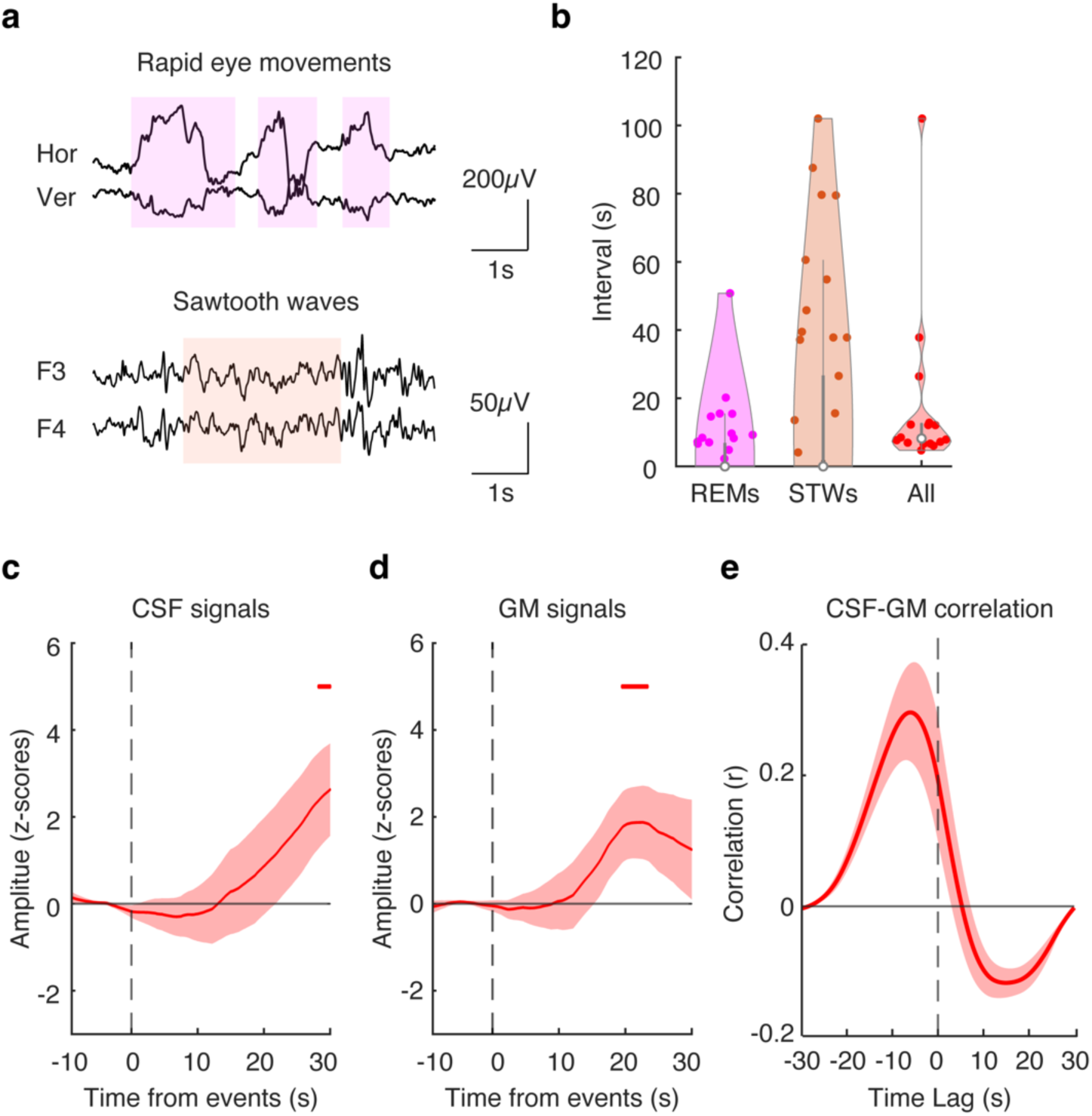
REM sleep events induce changes in CSF dynamics. **a**, Examples of horizontal (Hor) and vertical (Ver) rapid eye movements (top) and sawtooth waves detected from the frontal electrodes (bottom). **b,** Intervals of rapid eye movements (REMs), sawtooth waves (STWs) and all REM sleep events (All). **c,** CSF signal changes time-locked to the onset of REM sleep events. The horizontal bars represent statistical significance against zero baselines according to two-tailed one-sample *t*-tests (FDR-corrected *P* < 0.05). **d,** GM signal changes time-locked to the onset of REM sleep events. The horizontal bars represent statistical significance against zero baselines according to two-tailed one-sample *t*-tests (FDR-corrected *P* <0.05). **e**, Cross-correlation between CSF and GM signal changes time-locked to all REM events including rapid eye movements and sawtooth waves (max |r| = 0.31 at lag = –6s).

**Figure 3.**
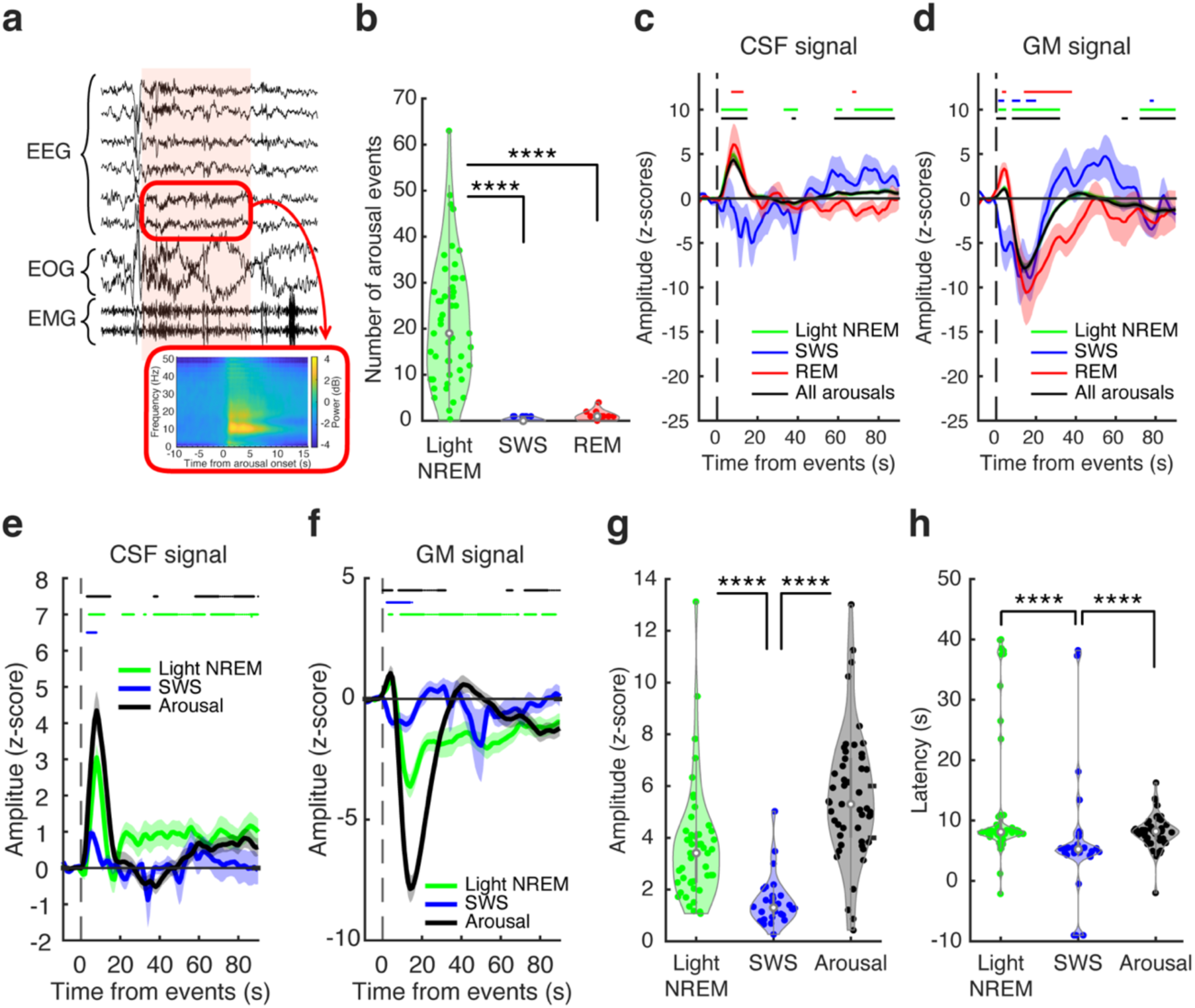
Arousals induce large CSF signal changes during sleep. **a**, Example of an epoch containing an arousal event (red). The embedded diagram shows an EEG spectrogram (O1 and O2 electrodes averaged) during arousal events (number of samples = 50 sleep sessions), indicating EEG activations across a wide range of frequency bands. **b,** Number of arousals during light NREM sleep, slow-wave sleep and REM sleep stages. Two-tailed Wilcoxson signed-rank tests with Bonferroni correction, \*\*\*\**Ps* < 0.001. **c,** CSF signal changes time-locked to arousal events for each sleep stage. The horizontal bars indicate significance against zero baselines via two-tailed one-sample *t*-tests (FDR-corrected *Ps* < 0.05). Because the majority of arousals occurred from light NREM sleep (as indicated in **b**), the plot for light NREM sleep (light green) and that for all arousal events (black) overlap. **d,** GM signal changes time-locked to arousal events. The horizontal bars indicate significance against zero baselines via two-tailed one-sample *t*-tests (FDR-corrected *Ps* < 0.05). As in **c**, because the majority of arousals occurred from light NREM sleep, the plot for light NREM sleep (light green) and that for all arousal events (black) overlap. **e,** CSF signal changes to arousal (black), slow waves during light NREM sleep (green, same as Fig. 1d), and slow waves during slow-wave sleep (blue, same as Fig. 1e). The horizontal bars indicate significance against zero baselines via two-tailed one-sample *t*-tests (FDR-corrected *Ps* < 0.05). **f,** GM signal changes to arousal (black), slow waves during light NREM sleep (green), and slow waves during slow-wave sleep (blue). The horizontal bars indicate significance against zero baselines via two-tailed one-sample *t*-tests (FDR-corrected *Ps* < 0.05). **g,** Peak amplitudes of CSF signals induced by slow waves during light NREM sleep (green), Bonferroni correction, \*\*\*\**Ps* < 0.001). The plots for light NREM and slow-wave sleep are the same as those in Fig. 1g. **h,** Mean latencies to reach the peak CSF signals induced by slow waves during light NREM sleep (green), slow-wave sleep (blue), and arousal (black). Two-tailed Wilcoxson signed-rank tests with Bonferroni correction, \*\*\*\**Ps* < 0.001. SWS, slow-wave sleep. The plots for light NREM and slow-wave sleep are the same as those in Fig. 1f.

### Different contributions of sleep brain oscillations and arousal mechanisms to CSF dynamics

Thus far, our results indicate that brain oscillations and neural events during deep sleep are tightly linked to CSF signal changes. Could these results be explained solely by cortical arousals, as arousals can occur throughout during sleep? If so, similar changes in the CSF signals could follow sleep-related brain oscillations and arousals. To investigate whether the contributions of sleep-related brain oscillations to CSF fluctuations are based on arousals, we examined arousal-associated CSF signal changes and directly compared them with slow wave-associated CSF signal changes. Arousal events in the EEG signals were detected by sleep experts in accordance with the standard criteria (29). The arousal events were defined as an abrupt shift in EEG frequency, which was clearly visible in the EEG signals as in the standard AASM criteria (**Fig. 3a**; ***Sleep event detection*** in **Materials and Methods**). The CSF and GM signals were subsequently segmented on the basis of each arousal onset.

The arousals occurred significantly more often during light NREM sleep (21.2 ± 1.9) than during slow-wave sleep (0.3 ± 0.1; **Fig. 3b**; Wilcoxson signed-rank test, two-sided with Bonferroni correction: *n* = 30, *z* = 4.7832, *P* = 1.7257e-06) or REM sleep (0.9 ± 0.2; **Fig. 3b**; Wilcoxson signed-rank test, two-sided with Bonferroni correction: *n* = 19, *z* = 3.8242, *P* = 1.3121e-04). These findings confirm that deep sleep is the least prone to arousals. Next, CSF signal changes time-locked to arousal events were measured. The average CSF signals increased 8 s after the onset of the arousal events and then returned to baseline 20 s after onset (**Fig. 3c**, black). Arousal-related CSF and GM signal changes were similar across sleep stages (**Fig. 3c, d**), with the exception that the CSF signal changes to arousals were barely visible during slow-wave sleep (**Fig. 3c**). Notably, the CSF and GM signal changes that followed arousals and slow waves during light NREM sleep had similar signal properties (**Fig. 3e, f**). The latency to reach the peak CSF signal was approximately 8 s (**Fig. 3e, h**), and CSF signal changes were followed by a large GM undershoot (**Fig. 3f**). In contrast, slow waves during slow-wave sleep had a shorter latency (**Fig. 3e, h**; Wilcoxson signed-rank test, two-sided with Bonferroni correction: *n* = 28, *z* = 3.3704, *P* = 7.5062e-04) and smaller amplitude changes (**Fig. 3e, g**; Wilcoxson signed-rank test, two-sided with Bonferroni correction: *n* = 28, *z* = 4.2810, *P* = 1.8603e-05) in the CSF signals as compared to the arousals (see **Fig. S5** for each comparison). One may wonder whether the differences in the signal properties between arousals and slow waves are simply due to the frequencies of the events, as a larger number of events may induce larger accumulative changes. However, since the total number of slow waves (22.7 ± 1.9; Wilcoxson signed-rank test, two-sided with Bonferroni correction: *n* = 28, *z* = 3.3933, *P* = 6.9063e-04), this is unlikely.

These findings consistently indicate that arousals and slow waves during light sleep are followed by slow and large CSF fluctuations, whereas slow waves during slow-wave sleep accompany fast yet small CSF signal changes, suggesting that sleep depth-dependent brain oscillations and arousals contribute differently to the CSF fluctuations.

### Faster frequency CSF signal components during slow-wave sleep and REM sleep

Our results thus far indicate that CSF signal changes are time-locked to brain oscillations and neural events independent of arousals. Furthermore, our results suggest CSF signals fluctuate *faster* during slow-wave sleep than during light sleep. If so, this should result in stronger faster frequency component in CSF-signal based power spectrum (38). To test this, we examined the periodic oscillation components of CSF signals across different sleep depths (see ***Power spectrum analysis of fMRI data*** in **Materials and Methods**; **Fig. S6**). Our results revealed that the faster component power (approximately 0.06-0.12 Hz) in the CSF signals was significantly greater during slow-wave sleep than during wakefulness or light NREM sleep (**Fig. S6a**, blue). In contrast, the slow component power (<0.06 Hz) in the CSF signals was significantly greater during light NREM sleep than during wakefulness or slow-wave sleep (**Fig. S6a**, green). CSF dynamics during REM sleep were characterized by a mixture of slow and very fast (>0.12Hz) components (**Fig. S6a**, red). Similarly to light NREM sleep, the slow component power was significantly greater during REM sleep than during slow-wave sleep or wakefulness. However, the power of the very fast component was significantly greater during REM sleep than during light NREM sleep (**Fig. S6a**). These findings demonstrate that both slow-wave sleep and REM sleep are indeed characterized by higher frequency CSF dynamics than light sleep.

### CSF signal changes associated with respiration, cardiac rhythms, and head motions

Since CSF signals are known to couple with respiration and cardiac pulses (39–41), we performed control analyses and confirmed that none of the respiratory or cardiac rhythms were likely to have caused the differences in the CSF signal changes among sleep stages and brain oscillations in this study. First, the intervals of the events were longer during the deep sleep stages than during the light sleep stages. Cardiac cycles were significantly longer during slow-wave sleep or REM sleep than during wakefulness (slow-wave sleep vs. wakefulness: Wilcoxson signed-rank test, two-sided with Bonferroni correction: *n* = 25, *z* = 4.3747, *P* = 1.2157e-05; REM sleep vs. wakefulness: Wilcoxson signed-rank test, two-sided with Bonferroni correction: *n* = 15, *z* = 3.3004, *P* = 9.7656e-04; **Fig. S7a**). There was no significant difference in respiratory cycles across different sleep depths (**Fig. S7b**). Second, the periodic frequency changes across different sleep depths were not produced by denoising processes during the fMRI preprocessing. To this end, we investigated the periodic component of CSF signals across different sleep depths without denoising processes of respiration and cardiac pulses following previous studies (25, 26). The key findings in the periodic frequency changes remained regardless of the existence of respiration and cardiac cycles in the fMRI signals (**Figs. S8**). Third, although a gradual increase occurred in the CSF signals after the respiration and cardiac pulses, these changes were significantly smaller than the CSF signal changes that occurred after brain oscillations (**Fig. S9**). Fourth, the cardiac and respirations had higher frequency signals (cardiac: around 1Hz; respiration: around 0.3Hz; **Fig. S10**) that did not overlap with the frequency range induced by sleep brain oscillations (<0.15 Hz). Additional control experiments and analyses were conducted to confirm that the aliasing noises from the rhythm of respirations and cardiac pulses (42) were not the main factor inducing the frequency peak observed at ∼0.06Hz in the CSF signals (**Fig. S11; *The control experiment and analyses to test aliasing noises*** in **Materials and Methods**). Thus, these physiological signals are not the main causes of the faster oscillating CSF signals found during slow-wave sleep and REM sleep.

Additional control analyses on head motions indicated that head motions were unlikely to have examined differences in the estimated distance of head motion during fMRI scans for each of sleep stages (see ***MRI data analysis*** in **Materials and Methods** for head motion measurement). We found that head motion during slow-wave sleep (0.021 ± 0.002 mm/volume) was significantly smaller than that during light NREM sleep (0.027 ± 0.002 mm/volume, Wilcoxson signed-rank test, two-sided with Bonferroni correction: *n* = 29, *z* = 4.7030, *P* = 2.5631e-06) or wakefulness (0.025 ± 0.002 mm/volume, Wilcoxson signed-rank test, two-sided with Bonferroni correction: *n* = 20, *z* = 3.9199, *P* = 8.8575e-05). Thus, the effect of head motion, if any, would have been minimal during deep sleep compared with light sleep. Next, correlation coefficients between head motion values and CSF and GM signals were measured by Pearson’s correlation. The group mean correlation coefficients were 0.06±0.02 (mean ± SE) between CSF signals and head motions and –0.09±0.03 (mean ± SE) between GM signals and head motions, indicating very small correlation with head motions.

### Relationship between CSF signal in the lateral ventricles and other regions

How might the CSF signals in the lateral ventricles relate to signals in the other regions? The present study extracted CSF signals from the lateral ventricles instead of the fourth ventricles. Thus, one might wonder whether these signals represent the ventricle signal itself or residuals produced by some artifacts (head motion artifacts). To check the relationship between the lateral ventricle signals and fMRI signals in other regions, we first examined zero-lag correlations between the CSF signals from the lateral ventricles and signals in each of the whole-brain voxels (see ***Voxel-wise correlation analyses: zero-lag correlation and cross-correlation time-lag map*** in **Materials and Methods**). In the group mean voxel-wise zero-lag correlation map (**Fig. S12a**), we found positive correlations within the ventricular regions, including the lateral and the fourth ventricles. We also observed wide-spread negative correlations between the lateral ventricle signals and grey matter regions. Importantly, the strongest positive correlations (*r* = 0.3) concentrate in the lateral ventricles.

Next, we examined the time-lag of the strongest positive correlation between the lateral ventricle signals and signals in each of the whole-brain voxels (see ***Voxel-wise correlation analyses: zero-lag correlation and cross-correlation time-lag map*** in **Materials and Methods**). We found that signals in the grey matter regions generally precede lateral ventricle signals, whereas small lags can be found within in the ventricular regions (**Fig. S12b**). Additionally, positive lags are found in the subarachnoid space, meaning the lateral ventricle signals precede the subarchnoid signal changes. These findings are consistent with a previous study investigated the fourth ventricle signals (43). Thus, the CSF signals in the present study represent ventricle signals, having tight temporal relationship with grey matter and subarchnoid signal changes.

### Sleep-depth-dependent and arousal-associated changes in the brain network

Because brain oscillations during slow-wave sleep are involved in plasticity and homeostatic regulation (44–46), while K-complexes during light sleep can accompany arousals (16, 17, 33, 34), we next asked whether the different modulations of CSF and GM signals found during light sleep, deep sleep, and arousals could be associated with different brain circuits involved during different depths of sleep. To this end, we applied a generalized linear model (GLM) analysis on the fMRI signals to each event. We found that different cortical networks were recruited depending on the depth of sleep (**Fig. 4**, **Tables S3–9**). The prefrontal and hippocampal regions, striatum, and amygdala were more strongly recruited to slow waves during slow-wave sleep than during light NREM sleep, consistent with previous studies investigating high-amplitude slow waves during slow-wave sleep (31) (**Fig. 4**, **Table S4**), whereas sensorimotor areas, including the parietal and visual cortices and cerebellum, were more strongly activated with slow waves during light NREM sleep than during slow-wave sleep, again replicating previous studies (47) (**Fig. 4**, **Table S3**). These findings indicate that different cortical processing underlies slow waves during light NREM vs. slow-wave sleep. Additionally, we found that more localized cortical brain regions were recruited to sleep spindles during slow-wave sleep than during light NREM sleep (48, 49) (**Tables S7 & S8**). the striatum, and the amygdala, also consistent with previous studies using fMRI and PET (50–52) (**Fig. 4, Tables S5 & S9**). In contrast to slow-wave sleep or REM sleep, arousals involved whole-brain activations not localized to any regions (**Fig. 4**, **Table S6**). Taken together, these results demonstrate that brain oscillations involved in regulating CSF dynamics recruit essentially different networks depending on the depth of sleep.

**Figure 4.**
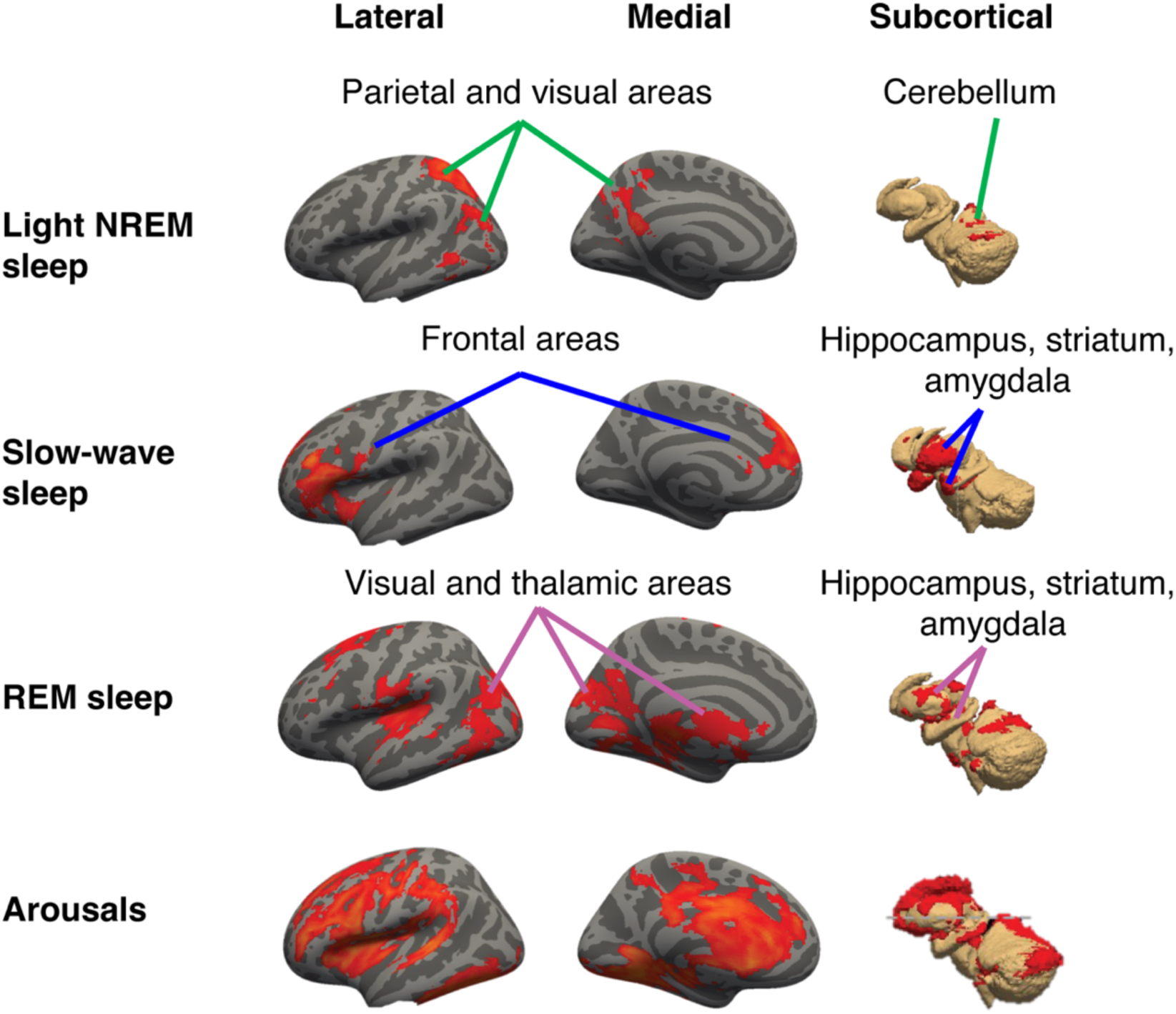
Sensory and motor regions are recruited during light sleep, whereas plasticity and homeostatic circuits dominate deep sleep. A generalized linear model (GLM) analysis was performed on the fMRI data corresponding to each event. A cluster-based statistical analysis was conducted at the group level, with the cluster-defining threshold (*Z* > 3.1) corrected at *P* < 0.05. Left, lateral view of the left hemisphere of a cortical surface in an inflated format. Middle, medial view of the left hemisphere of a cortical surface in an inflated format. Right, subcortical areas. The red highlighted areas in the first row indicate the areas that significantly increased during slow waves in light NREM sleep compared with those in slow-wave sleep. The red highlighted areas in the second row indicate the areas that significantly increased during slow waves in slow-wave sleep compared with those in light NREM sleep. The red highlighted areas in the third row indicate the areas that increased significantly with rapid eye movements (REMs) during REM sleep. The red highlighted areas in the fourth row indicate the areas that increased significantly in response to arousals. Light NREM sleep induces sensory and motor regions, whereas slow-wave sleep induces regions involved in plasticity and homeostatic circuits. REM sleep recruits visual and plasticity circuits. Arousals accompany widespread brain activation. Details of brain regions recruited for each sleep event can be found in **Supplementary Tables S3-9.**

## Discussion

The present results robustly indicate that deep human sleep has a unique way of interacting with CSF signals, as indicated in **Table S2**. Slow waves and sleep spindles during slow-wave sleep are followed by faster changes in CSF fluctuations. Neural events during REM sleep, such as rapid eye movements and sawtooth waves, also accompany CSF signal changes. We further revealed distinct frequency profiles in CSF signals among different sleep stages. The frequency of CSF signals become faster during deep sleep than during wakefulness or light sleep. In contrast, arousals and brain oscillations during light sleep were followed by infrequent, sharp CSF changes. We confirmed that different brain networks are activated after brain oscillations or events during different depths of sleep. Deep sleep recruits brain regions involved in memory and homeostatic regulation, whereas light sleep and arousal recruit whole-brain activation, including the sensory-motor network.

The present study suggests that at least two different processes are involved in regulating CSF dynamics in human sleep. The first process is prominent during light sleep when arousals and arousal-related brain activities, such as K-complexes, take the lead in modulating CSF oscillations. The second process is facilitated specifically during deep stable sleep when arousals are largely suppressed. These results further suggest that during deep sleep, CSF oscillations are facilitated by spontaneous brain oscillations and neural events, such as slow waves, sleep spindles, and REM sleep events. Since sleep events occur more often than arousals during deep sleep, it is highly likely that sleep events, not arousals, are the major contributors to CSF oscillations during deep sleep. We propose that these two processes are supported by distinctive mechanisms, as shown in **Fig. 5**. The process during light NREM sleep is based on arousal mechanisms (“arousal-based mechanism”), in which transient increases in brain activity facilitate CSF oscillations by arousing stimuli to subcortical regions. Another process that we propose here functions specifically during deep sleep (“deep sleep-based mechanism”), which is tightly linked to sleep brain oscillations to drive the CSF oscillations. This deep-sleep mechanism may work continuously, producing faster CSF oscillations by tightly interacting with sleep-related brain oscillations. In contrast to the arousal mechanism that induces sharp and transient CSF signal increase, the deep-sleep mechanism may induce low-amplitude oscillations, potentially causing minimal load to the vascular system while enabling continuous faster oscillations.

**Figure 5.**
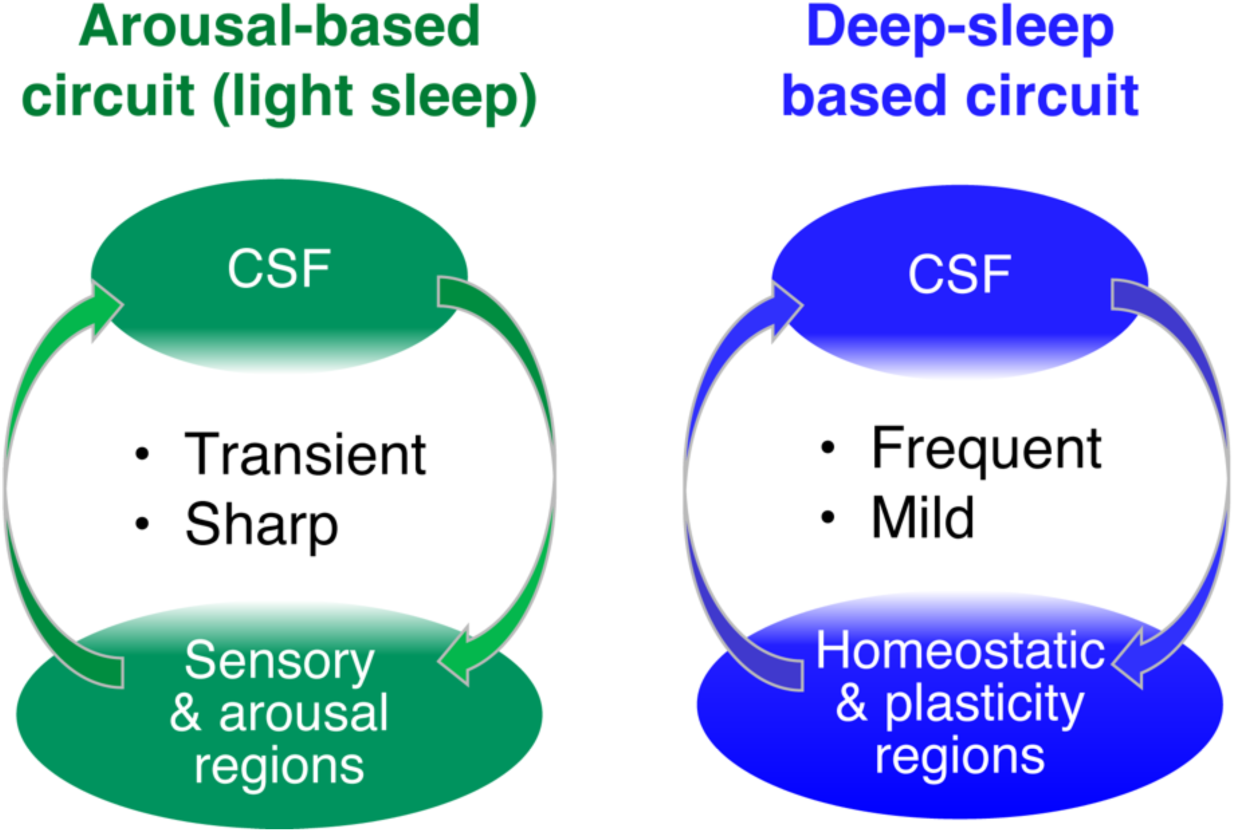
An illustration of the hypothetical relationships between CSF and brain circuits during light and deep human sleep. During light sleep (left), CSF oscillations are characterized by intermittent sharp peaks induced by cortical arousals or arousal-related waves (K-complexes). During deep sleep (right), CSF oscillations become faster, but their amplitudes are low (mild). These faster CSF oscillations are induced by sleep-related brain oscillations.

Although the direct relationship between CSF signal changes during deep sleep and waste clearance is unclear from the present study, the present results provide some important implications. First, we found that slow waves are among the main oscillations that produce faster CSF circulation during deep sleep. Given the tight relationship between slow waves and homeostatic sleep needs (7, 8, 44), faster CSF circulation associated with slow waves might work well for clearing waste products that might have accumulated during prolonged wakefulness. Second, the present results suggest that sleep spindles also accelerate CSF oscillations. Sleep spindles are known to facilitate episodic and motor learning through plastic changes in cortical and subcortical regions (53–55). Our analysis has indeed shown that crucial brain network regions involved in episodic and motor learning (e.g., hippocampus and striatum) were recruited during sleep spindles during slow-wave sleep, consistent with previous studies (48). Thus, it is possible that CSF oscillations are coupled not only to sleep homeostasis but also to plasticity processes, where sleep spindles might play a major role.

While we found faster CSF dynamics during REM sleep as well as during slow-wave sleep, the roles of CSF oscillations might differ. First, the latency to the CSF peak was much longer during REM sleep than during slow-wave sleep. Second, we found that not only the very fast component (> 0.12 Hz) but also the slow component (< 0.06 Hz) was present in the CSF frequency profiles during REM sleep. Third, the CSF peak did not accompany the GM undershoot during REM sleep. In addition to the rapid eye movements and sawtooth waves we analyzed in this study, EEG theta and alpha activities can be observed, and muscle twitches occur intermittently during REM sleep, which were not examined here. Since sympathetic nervous system activity, which regulates heart rates and blood pressure, is increased during REM sleep, it is possible that autonomic variability (e.g. respiratory and cardiac variabilities) in REM sleep also contributes to the CSF signal changes during REM sleep, as was proposed previously (39, 56).

One may wonder whether cardiac and respiratory rhythms cause differences in the CSF peaks during different sleep stages and brain oscillations since CSF signals are known to be coupled to cardiac and respiratory rhythms (40, 57, 58). However, these signals are unlikely to be the sources of the differences in the CSF signal changes among sleep stages and brain oscillations found in the present study for the following three reasons. First, the intervals of the cardiac signals were significantly longer during slow-wave sleep or REM sleep than during wakefulness. There was no significant difference in the respiratory signals across different sleep depths. If these factors contribute to changes in the CSF signal, the CSF signals should be slower during deep sleep than during light sleep. Thus, these rhythms are not in accordance with the significantly faster CSF signals found during slow-wave sleep and REM sleep. Second, we investigated the effects of respiration and cardiac pulses in fMRI signals on the periodic frequency changes in CSF signals. The key findings of the periodic frequency changes in the CSF signals remained regardless of the existence of respiration and cardiac pulses in the fMRI signals. Third, previous studies have indicated that cardiac– and respiration-driven CSF signal changes occur 0.45–1.58 s after the events (40). However, CSF signal changes caused by slow waves were in the range of 5.5–8 s after the events, much later than the cardiac and respiration-driven changes that might have occurred. Additionally, we investigated the CSF signal changes that followed respiration and cardiac pulses. We found that respiration– and cardiac-driven signal changes were significantly smaller than sleep-related brain oscillations and arousals. Thus, the CSF peaks caused by cardiac and respiratory rhythms, if any, should not overlap with the CSF peaks that appeared after brain oscillations. Finally, the results of control analyses showed that the aliasing noises from respirations and cardiac pulses were unlikely to be the main factor inducing the frequency peak observed at ∼0.06Hz in the CSF signals found in the present study.

The present study at least partially suggests why and how deep sleep matters for healthy human cognitive functions. Sleep disturbances are a major issue worldwide and are associated with various health-related issues, including cognitive impairments, neurodegenerative diseases and aging (59–62). Importantly, the most striking change found in these cases is the reduction in deep stable sleep. In other words, light NREM sleep most likely remains regardless of the severity of the disorder. Along with evidence showing a strong relationship between the stages of sleep, CSF dynamics, and metabolite clearance (5, 7, 8, 63), dysfunctions in deep sleep may have severe impacts on cognitive functions at least partially via altered CSF flows.

Although the present study did not directly measure the underlying mechanisms of the tight link between sleep brain oscillations and the CSF signals, one of the likely mechanisms may involve vascular dynamics. The vascular dynamics may change the size of the perivascular spaces of the penetrating arterioles where the CSF circulate throughout the brain (8, 64). Since neural activity, vascular dynamics, and CSF dynamics tightly interact with each other (11, 65), synchronous neural activities, such as slow waves, sleep spindles, and various REM-sleep events may trigger vasomotion, which may promote CSF circulation. Based on the present findings, which showed that deep sleep involves accelerated CSF signal changes, these underlying processes may be suggest that deep sleep has a specific role in maintaining the tight physiological and neural balance and that light sleep may not be able to compensate for the loss of deep sleep caused by sleep restrictions, aging, or diseases.

The following four points need to be addressed in future research. First, our data were obtained from midafternoon naps. Thus, what happens during nightly sleep when sleep needs are greater is unclear. As the duration of slow-wave sleep is the longest and the number of slow waves is the maximum during the first half of nightly sleep (29), it is possible that CSF oscillations are much greater. REM sleep, in contrast, is longest during the early morning, and REM density is greater (29). Thus, REM-related CSF dynamics could benefit from early morning studies. Additionally, our data are restricted to approximately 1.5 hours, corresponding to the first sleep cycle. Thus, how CSF signal fluctuations change over multiple sleep cycles is unclear. Second, the present study used a sparse fMRI sequence with longer TR (3s) than previously used (∼1 s) to prioritize the acquisition of deep sleep data, rather than using a fast fMRI method to measure the inflow effects in the CSF signals from the fourth ventricle (11, 66, 67). Nevertheless, consistent with the previous studies investigated the lateral ventricles (68–70), our data clearly indicate that CSF signals are differently modulated during wakefulness vs. sleep. Third, it was not optimized to measure the signal changes driven by autonomic activity, including the cardiac and respiratory rhythms that were associated with CSF pulsations (12, 13, 39, 70). Fourth, since the present study did not directly measure the amount of metabolite waste or investigate the causal relationship between CSF and metabolites, how CSF during deep sleep contributes to waste clearance in humans needs to be clarified in future research.

Taken together, the results of the present study reveal that CSF signal changes are tightly associated with sleep-brain oscillations and events during slow-wave sleep and REM sleep. These CSF dynamics during deep sleep could be distinct from the arousal-based CSF dynamics for the following reasons. First, the latency and amplitude characteristics are different. Second, the frequency components are different among different sleep stages. Third, the relationships with whole GM signals are different. Fourth, the brain network changes after sleep brain oscillations during light vs. deep sleep and after arousals are different. While the causal relationship between CSF signals during sleep and brain functions is unclear, the present study collectively suggests that deep sleep-based CSF dynamics are the key to understanding how healthy brain functions are maintained through sleep in humans.

## Materials and Methods

### Participants

Twenty-five young healthy participants (mean age of 23.6 ± 4.8 years, 15 female) participated in this study. All of the participants provided written informed consent for their participation prior to the start of the study. This study was approved by the institutional review board at RIKEN. There were no studies that tested CSF dynamics during deep sleep prior to the present study. Previous studies using fMRI during sleep in human participants (11, 47, 71) have reported n=11–20 subjects. However, to ensure that the results of our study were reliable and replicable, we used 25 subjects.

### Screening process for eligibility

Before consent was obtained, the screening process for eligibility took place to confirm the following aspects. First, the participants were between 18 and 35 years old. Second, on the basis of a self-report questionnaire (23), anyone who had neurological or psychiatric conditions (e.g., epilepsy, migraine, stroke, chronic pain, major depression, anxiety disorder, psychotic disorder), was currently using medication, or was suspected of having a sleep disorder (including insomnia, sleep apnea syndrome, hypersomnolence, restless legs syndrome, parasomnias) was not eligible to participate. Third, participants who had an irregular sleep schedule, i.e., those whose eligible. We also ensured that the subjects had not taken a trip to a different time zone during the last six months prior to the experiment. Participants who worked night shifts or who had a habit of taking a nap were not eligible for this study. Fourth, a Pittsburgh Sleep Quality Index (PSQI) was used to assess sleep quality and disturbances over a one-month time interval before the experiments, and a PSQI score within 5 was used as an inclusion criterion (72). Fifth, those who failed to complete an MRI safety questionnaire (i.e., claustrophobic, pregnant, having metallic objects in the body, such as vascular clips, prosthetic valves, metal prostheses, metal fragments, pacemakers, or dental braces) were not eligible for this study.

### Experimental design

All eligible subjects participated in two sleep sessions approximately one week apart. The participants arrived at the experimental room at approximately noon. The participants first answered a questionnaire to report their bedtime, wake-up time, and duration of sleep the night before. They also answered the Stanford Sleepiness Scale (SSS) rating (73), which ranges from 1 (feeling active, vital, alert, or wide awake) to 7 (no longer fighting sleep, sleep onset soon; having dream-like thoughts). The subjects were asked to choose the scale rating that described their state of sleepiness. Then, the electrodes for PSG were attached, which took approximately one hour (see the PSG measurement section below). After the electrodes were attached, each sleep session started in the early afternoon for simultaneous measurement of MRI and PSG (see the MRI acquisition section below). The sleep session was conducted for less than two hours in the MRI scanner including structural and functional image acquisition. After the sleep session, they answered a questionnaire about the sleep content (data not shown).

The participants were instructed to maintain their regular sleep-wake habits, i.e., their daily wake/sleep time and sleep duration, until the study was over. The sleep-wake habits of the participants were monitored by a sleep log and an actigraphy device (MTN-221, ACOS, CO., LTD, Japan; SleepSign Act ver.2, KISSEI COMTEC, CO., LTD, Japan) during the study. Throughout their participation, the subjects were instructed to refrain from taking naps. One day prior to the experiments, alcohol consumption was not allowed. Unusual excessive physical exercise and caffeine consumption were not allowed on the day of the sleep sessions.

### MRI acquisition

MRI data were acquired with a 3T Siemens Prisma MRI scanner (Siemens Healthineers, Germany) using a 64-channel head and neck coil at RIKEN. An anatomical T1-weighted image was acquired using a 3D magnetization-prepared rapid gradient echo (MPRAGE) sequence (TR = 2180 ms, TI = 1100 ms, TE = 2.95 ms, FA = 8°, voxel size = 0.7×0.7×0.7 mm^3^, FOV = 290 × 320 × 256 mm). Functional MRI data were acquired using a sparse gradient-echo echo-planar imaging sequence (TR/TE=3000/30 ms, FA=85°, Multiband factor=3, 1800 volumes, GRAPPA factor=3, bandwidth=752 Hz/pixel, B1+RMS=0.5 μT, MR acquisition time=0.9 s, and quiet period=2.1 s) (74). 42 transverse slices (voxel size = 3×3×3 mm^3^) orientated parallel to the AC-PC plane were acquired if successfully covering the entire cerebrum. If not, the slice alignments were adjusted accordingly to cover the entire cerebrum. Cardiac and respiratory cycles were simultaneously recorded using a fingertip PPG sensor and respiratory belt (BIOPAC, US) with a 100 Hz sampling rate. Anatomical and functional MRI data were acquired simultaneously with polysomnography (see the ***PSG measurement*** section below). Cushions and gauze were used to stabilize the subjects’ heads to reduce discomfort and head motion. Back and knee cushions were used upon the participants’ request to further reduce discomfort. Several blankets were used to keep the participants warm and to initiate sleep during the scan (22, 23).

### PSG measurement

EEG data were acquired using a BrainAmp MRplus EEG amplifier (Brain Products, Gilching, Germany) with a 5 kHz sampling rate and an MR-compatible 32-channel EEG cap (EasyCap, Wörthsee, Germany). The hardware bandpass filters were set to a 0.016–250 Hz range, with a international 10-20 system with additional drop-down channels or bipolar electrodes for recording the electrocardiogram (ECG), electrooculogram (EOG), and chin electromyogram (EMG). EEG was measured from either 27 electrodes (16 participants) or 31 electrodes (9 participants). For all of the participants, EOG was measured from electrodes placed at the outer canthi. EMG was measured from the mentum. ECG signals were measured from the lower shoulder blade. FCz was used as the reference electrode, whereas AFz was used as the ground electrode. EEG data were transferred outside the scanner room through fiber optic cables to a computer running the BrainVision Recorder software (Brain Products, Gilching, Germany). MR-EEG scanner clocks were synchronized (Brain Products Synchbox) for all EEG data acquisitions, and the onset of every scanner repetition time (TR) period was recorded in the EEG data for MR gradient artifact correction. The impedance of all electrodes was initially maintained below 10 kΩ and was maintained below a maximum of 20 kΩ for the duration of the study.

### PSG data preprocessing

PSG data were contaminated by artifacts due to the strong magnetic field. To remove the artifacts from the PSG data, gradient artifacts were first corrected using BrainVision Analyzer2 (Version 2.2.0, Brain Products GmbH, Gilching, Germany) with the standard average artifact subtraction (AAS) method. The AAS method uses sliding window templates formed from the averages of 21 TRs, which are subtracted from each occurrence of the respective artifacts for each electrode (75, 76). The gradient-artifact corrected data were subsequently bandpass filtered (EEG: 0.1–50 Hz with a notch filter of 50 Hz) and downsampled to 250 Hz. Following the GA correction, cardiac R-peaks were automatically detected from the ECG recording in BrainVision Analyzer2, checked visually and corrected manually if necessary. These R-peak events were used to inform ballistocardiogram correction. The BCG artifacts were further corrected via the AAS method with sliding window templates formed from the averages of 21 R-peaks in BrainVision Analyzer2 (10, 77). These procedures were conducted for the whole duration of the fMRI scan to obtain artifact-removed PSG data for further processing (see ***Sleep stage scoring*** and ***Sleep event detection*** below).

### Sleep stage scoring

Sleep staging was conducted on the artifact-removed PSG data following the standard criteria (29). Prior to sleep staging, PSG data were further bandpass filtered between 0.3 and 40 Hz for EEG, EOG and ECG and between 10 and 50 Hz for EMG to remove low-frequency drift and high-frequency noise and were re-referenced to the linked mastoids (TP9 & TP10). Scoring of the sleep stages was performed every 30 s in conjunction by two trained sleep scorers with over twenty years of experiences (MT & AS). We obtained sleep parameters, including the total sleep time and duration of stage wakefulness (W), NREM 1 (N1), NREM 2 (N2), NREM 3 (N3), and stage REM (REM) (**Table S1**). For further analysis, N1 and N2 were combined as light NREM sleep, and N3 was categorized as slow-wave sleep. Epochs with motion artifacts and arousal events (29) were marked and removed from further analyses, whereas the arousal events were used only for arousal analysis. Motion artifacts were defined as an epoch containing artifacts that lasted longer than 1s or a participant showing motion in the video recording.

### Sleep event detection

Using the artifact-removed PSG data, sleep events, including slow waves, sleep spindles, rapid eye movements, sawtooth waves, and arousal events, were detected as follows.

#### Slow waves

Slow waves were detected semiautomatically as follows. First, EEG signals were bandpass filtered between 0.1 and 4 Hz with a zero-phase 4th-order Butterworth filter. Then, slow waves were detected automatically in accordance with a previous study (30) using Wonambi software (https://wonambi-python.github.io/) from the F3, Fz, F4, C3, Cz, C4, P3, Pz, and P4 electrodes during the N2 and N3 stages. Only the waves that met the following three criteria were selected and a subsequent negative-to-positive zero crossing separated between 0.3 s and 1 s, 2) a negative peak between the two zero crossings with voltages smaller than –80 μV, and 3) a peak-to-peak amplitude greater than 140 μV. The automatically detected slow waves were then reviewed and removed by an experimenter if necessary. The number of slow waves was measured for each electrode, and the electrode that had the maximum number of slow waves was used as the most sensitive electrode for slow wave detection. The onset and duration of slow waves were measured from the specific electrode for further analysis of CSF and GM signals (see ***fMRI time course analysis*** and ***fMRI GLM analysis*** below).

#### Sleep spindles

Spindles were detected semiautomatically. As the first step, spindles were detected automatically (32) using Wonambi software from the F3, Fz, F4, C3, Cz, C4, P3, Pz, and P4 electrodes during the N2 and N3 stages in the following steps: EEG signals were filtered between 12–15 Hz, which corresponded to the sleep spindle frequency band. Then, the root mean square (RMS) of the filtered signals was subsequently calculated at every sample point using a moving window of 0.2 s. The threshold for spindle detection in the RMS signal was set to 1.5 standard deviations of the filtered signal for each electrode. A spindle was detected when the RMS signal remained above the threshold for 0.5–3 s, and then the beginning and end of the spindle were marked at the threshold crossing points. When sleep spindles from multiple electrodes overlapped within a 5 s time window, those sleep spindles were treated as trains of continuous sleep spindles and merged. In the second step, the automatically detected spindles were reviewed and removed by an experimenter if necessary. The onset and duration of sleep spindles were used for further analysis of CSF and GM signals (see ***fMRI time course analysis*** and ***fMRI GLM analysis*** below).

#### Rapid eye movements

Rapid eye movement events were detected during REM sleep. Rapid eye movement events were automatically detected on two EOG signals (78). Specifically, to separate rapid eye movements from slow eye movements (SEMs), two channels of EOG data are filtered using a fourth-order Butterworth bandpass filter with cutoffs at 1 and 9 Hz to yield the filtered EOG data. The candidate REM sleep events were generated as the negative instantaneous product of the two filtered EOG sequences, and the REM sleep events were detected on the basis of their amplitude threshold. The amplitude threshold was arbitrarily determined on the basis of each individual dataset, since, owing to differences in the setup and/or the quality of the artifact removal, there were individual differences in the amplitude of the EOG data. The onset and duration of rapid eye movements were used for further analysis of the CSF and GM signals (see ***fMRI time course analysis*** below).

#### Sawtooth waves

Sawtooth waves were manually marked on the scalp EEG independently by two trained sleep scores (MT & SA) for the REM sleep episodes. These waves were defined as bursts of consecutive surface-positive 2–5 Hz frontocentral bilateral synchronous symmetric waves with amplitudes of 20–100 µV and with a slow incline to a negative peak with a subsequent steep linear decline ending at a positive peak (79–81). The minimum number of consecutive waves was set as 3. The consensus of the markings obtained from the two raters was selected, and the onset and duration of sawtooth waves were used for further analysis of the CSF and GM signals (see ***fMRI timecourse analysis*** below).

#### Arousals

An arousal was defined as an abrupt shift in EEG frequency, which may include theta, alpha and/or frequencies greater than 16 Hz (but not spindles) that were at least 3 seconds in duration, with at least 10 seconds of stable sleep preceding the change, in accordance with the standard AASM criteria (29). Arousal events were manually scored by two trained sleep experts (MT & SA) onset and duration of arousal were used for further analysis of the CSF and GM signals (see ***fMRI time course analysis*** and ***fMRI GLM analysis below)*.**

### MRI data analysis

High-resolution T1-weighted images were processed using FreeSurfer 7.4.0 (82) (https://surfer.nmr.mgh.harvard.edu/fswiki/FreeSurferWiki) to segment the head into five tissues (i.e., the scalp, skull, cerebrospinal fluid (CSF), gray matter and white matter), and an anatomical atlas. Anatomical surfaces such as the pial surface, gray/white matter interface and mid surface (i.e., a middle layer of the gray matter) were estimated using FreeSurfer (83). The lateral ventricles and whole gray matter region (GM) were extracted as the two main regions of interest (ROIs) from each individual participant’s anatomical image. The fMRI data were motion-corrected and slice-timing corrected using the AFNI (https://afni.nimh.nih.gov) and RIKEN fMRI processing programs (http://www.fmri.brain.riken.jp/∼mri/Software.html). Furthermore, the fMRI data were denoised using cardiac and respiration signals (25, 26). The two ROI images (GM and lateral ventricles) were subsequently registered to each fMRI image using FSL FLIRT (https://fsl.fmrib.ox.ac.uk/fsl/fslwiki/FLIRT), and the fMRI signals were z score transformed and averaged across the voxels inside each ROI for each sleep session and each participant. The distances of head motion during fMRI scans were estimated by MCFLIRT in FSL (84, 85).

### fMRI time course analysis

fMRI data in each ROI were segmented based on sleep events (slow waves, spindles, rapid eye movements, sawtooth waves, and arousals). First, –9–0 s (3TRs) before the onset of the events was defined as a baseline. This baseline signal was averaged and subtracted from 0–30 s after the onset of the events. These data were averaged across the number of each event for each sleep session and then averaged across the sleep sessions and participants.

### Cardiac and respiration peak-related fMRI time course analysis

Cardiac and respiratory cycles during each sleep session were recorded using a fingertip PPG sensor and respiratory belt (BIOPAC, US) with a 100 Hz sampling rate. Peaks of the cardiac and respiration cycles were detected using RIKEN fMRI processing programs (http://www.fmri.brain.riken.jp/∼mri/Software.html). fMRI data in each ROI were segmented based on the peaks of the cardiac and respiration signals. First, –9–0 s (3TRs) before the onset of the peak events was defined as a baseline. This baseline signal was averaged and subtracted from 0–30 s after the onset of the peak events. These data were averaged across the number of each event for each sleep session and then averaged across the sleep sessions and participants.

### Power spectrum analysis of fMRI data

Time-frequency analysis was applied to the whole fMRI signals during a sleep session of the two ROIs using a multitaper approach (74, 86) in the Fieldtrip toolbox (87) (http://www.ru.nl/neuroimaging/fieldtrip). Time windows of 200 s duration were moved across the data in steps of 3 s corresponding to 1 TR, resulting in a frequency resolution of 0.005 Hz. The calculated time-frequency representations (TFRs) were then transformed to the scale of dB by 10*log10, then demeaned by subtracting the mean power at each frequency during the whole recording to calculate power fluctuations. Then, they were segmented based on sleep stages every 30 s and averaged across each 30 s to extract each epoch power spectrum after removing arousal– and motion-contaminated epochs. On the basis of the resulting power spectrum, periodic components of the power spectrum were extracted after the estimated 1/f aperiodic components were removed by fitting oscillations and one-over-f (FOOOF) algorithms (88). The resulting periodic power spectra were averaged for each sleep stage first and then across sleep sessions for each ROI.

### Correlation analysis between CSF and GM signals

Sleep-specific event-locked (i.e., slow waves, spindles, and REM sleep events) time courses of correlations between the CSF and GM signals. The correlation coefficients and time lags were extracted when the absolute coefficient values were at their maximum between the two signals.

### Voxel-wise correlation analyses: zero-lag correlation and cross-correlation time-lag map

To calculate voxel-wise zero-lag correlation, voxel-wise maps of correlation between the CSF signal from the lateral ventricles and all voxels were computed using Pearsons’ correlation separately for each nap session. Next, we applied a Fisher Z transform prior to averaging then group mean group mean voxel-wise zero-lag correlation map was generated by averaging the individual maps. We also performed two-tailed one-sample t-tests (uncorrected P <0.001) to identify voxel regions with significant differences in correlation with the CSF signal compared with zero.

To generate cross-correlation time-lag map, time-lag of the maximum positive correlation between the CSF signal from the lateral ventricles and all voxels were computed using cross-correlation separately for each nap session. Next, group mean group mean voxel-wise time-lag map was generated by averaging the individual maps.

### CSF peak locked EEG topography analysis

Positive peaks of the CSF signal from the lateral ventricles during each sleep session were detected using a MATLAB function (findpeaks.m). The MRI-related artifact-removed EEG data were segmented based on the positive peak events of the CSF signals (–10 s to 0 s). The segmented EEG data were averaged across the sleep sessions and participants for each sleep stage (N2 and N3 sleep stages). EEG topographies across time were generated in the Fieldtrip toolbox (87) (http://www.ru.nl/neuroimaging/fieldtrip).

### Power spectrum analysis of cardiac and respiration signals

Cardiac and respiratory cycles during each sleep session were recorded using a fingertip PPG sensor and respiratory belt (BIOPAC, US) with a 100 Hz sampling rate. The same power spectrum analysis steps were applied to the cardiac and respiration signals (see ***Power spectrum analysis of fMRI data*** above).

### fMRI GLM analysis

MRI data were processed using FSL v6.0.7.7 (https://fsl.fmrib.ox.ac.uk/fsl/). Data from each participant were motion corrected (MCFLIRT), spatially smoothed (5 mm FWHM Gaussian kernel), high-pass temporally filtered (100 s cutoff), registered to their T1 anatomical brain image (FLIRT), and normalized to the MNI 2 mm standard brain. GLM analyses were performed using FEAT v6.00. On the basis of each sleep-specific event (onset and duration for slow waves, sleep spindles, and arousals) detected from the simultaneous EEG data, event-specific first-level analysis was performed employing one regressor: 1) the event onset and duration. The regressor of interest was convolved with the double-gamma HRF. Motion parameters estimated during image realignment (3 translations and 3 rotations around the x, y, and z axes) were included in the matrix as confounding regressors of no interest. Additionally, since MRI BOLD signal amplitudes vary across different vigilance states (11, 89–92), the mean MRI signal inside the white matter region and the mean MRI signal in the cerebrospinal fluid region were included in the GLM as nuisance regressors (93). For each event of each participant, the event-specific first-level results were combined across sleep sessions using a second-level, fixed effect analysis to calculate an average response per subject. For each event, these second-level results were subsequently combined and analyzed at the group level using a FLAME mixed-effect analysis (94). In a cluster-based threshold statistical analysis, the resulting group-level BOLD Z statistic images were thresholded using Z > 3.1 and a cluster-corrected significance threshold of *P* < 0.05. For visualization, the group-level thresholded significant BOLD Z statistic image for each event (slow waves, sleep spindles, and arousals) was transformed onto a standard template surface map (fsaverage) using feat2surf and a standard template subcortical brain region, and the group-FreeSurfer freeview software. The automated anatomical atlas 3 (AAL3 (95)) was used for anatomical labeling.

### The control experiment and analyses to test aliasing noises

We conducted the additional experiment using TR=2.5s to check the effect of the aliasing noise from the respirations and cardiac pulses on the CSF signals (42). If the spectrum peaks of the CSF signal (around 0.06Hz) were due to the aliasing noise, the 0.06Hz peak should disappear when the data were acquired with TR=2.5s since the aliasing noise between MRI acquisition rate (0.4Hz) and respiration peak (around 0.27Hz) would be 0.13Hz (0.4-0.27=0.13Hz) as compared to 0.06Hz (0.33-0.27=0.06Hz) when using TR=3s (0.33Hz).

Three young healthy participants (mean age of 25.3 ± 4.0 years, 3 female) participated in the control experiment. All of the participants provided written informed consent for their participation prior to the start of the study. The experimental design and data analyses were the same as the main experiment except that the TR for MRI acquisition was 2.5s instead of 3s. Time-frequency analysis was applied to the whole fMRI signals during a sleep session of the two ROIs using a multitaper approach (see ***Power spectrum analysis of fMRI data*** in **Materials and Methods**). We confirmed that the major spectrum peak of the CSF signals was still visible at ∼0.06Hz (**Supplementary Fig. S9**). These indicate that this spectrum peak of the CSF signals is not due to the aliasing noises between the MRI acquisition rate and respiration peak.

## Statistical analysis

The alpha level was set as 0.05. All tests conducted in this study were two-tailed. For the fMRI power spectrum analysis, nonparametric cluster-based permutation tests were used to find significant differences (*P* < 0.05) in the periodic power spectrum across different sleep stages in each ROI using the Fieldtrip toolbox (96–98). For the fMRI time course analysis, two-tailed one-sample *t*-tests (*P* <0.05) were applied to each time point of the fMRI signals compared with zero. Two-tailed paired *t*-tests were used to compare each time point of event-locked fMRI signals between sleep stages. For the statistical inference of the correlation analysis and the numbers and time intervals of each event across different sleep depths, the Shapiro-Wilk test was conducted to test whether the data were normally distributed. If data were not normally distributed, the Wilcoxon signed-rank tests were used. When normality was not rejected by the Shapiro-Wilk test, two-tailed paired *t*-tests were used. To control for multiple comparison problems, Bonferroni corrections were applied to minimize the false-positive rates in the statistical inference of the correlation analysis and the numbers and time intervals of each event across different sleep depths. False discovery rate (FDR) corrections (99) were used for other statistical comparisons, such as comparing each time point of the fMRI signals and the fMRI power spectrum. We also included a 95% CI for a significant *t*-test and Pearson’s correlation analysis. Statistical tests were run by MATLAB (2020b, MathWorks, Inc.).

## Data availability

The datasets generated and/or analyzed in the current study are available from the corresponding author upon request.

## Code availability

No custom algorithm or software was used to that is central to the present findings. The computer codes that were used to generate the results of this study are available from the corresponding author upon request.

## Inclusion & Ethics

This study was approved by the institutional review board at RIKEN. All of the participants provided written informed consent for obtaining and sharing individual-level data.

## Acknowledgments

The authors thank M. A. Carskadon and H. Lau for helpful comments on an earlier draft. The authors also thank H. Nakatomi, M. Gomyo, T. Yoshioka, K. Fukushima, M. Koyanagi for fruitful discussions on CSF measurements. This work was supported by JSPS KAKENHI Grant Numbers JP22H01107 (MT) and JP22K18664 (MT), the Naito Foundation (MT), Takeda Science Foundation (MT), Life Science Foundation of JAPAN (MU).

## References

1. R. Stickgold, Sleep-dependent memory consolidation. Nature 437, 1272–1278 (2005).

2. L. Marshall, H. Helgadóttir, M. Mölle, J. Born, Boosting slow oscillations during sleep potentiates memory. Nature 444, 610–613 (2006).

3. A. J. Krause, et al., The sleep-deprived human brain. Nat Rev Neurosci 18, 404–418 (2017).

4. M. R. Irwin, M. V Vitiello, Implications of sleep disturbance and inflammation for Alzheimer’s disease dementia. Lancet Neurol 18, 296–306 (2019).

5. L. Xie, et al., Sleep Drives Metabolite Clearance from the Adult Brain. Science (1979) 342, 373–377 (2013).

6. J. J. Iliff, et al., A Paravascular Pathway Facilitates CSF Flow Through the Brain Parenchyma and the Clearance of Interstitial Solutes, Including Amyloid β. Sci Transl Med 4 (2012).

7. P. K. Eide, et al., Altered glymphatic enhancement of cerebrospinal fluid tracer in individuals with chronic poor sleep quality. Journal of Cerebral Blood Flow & Metabolism 42, 1676–1692 (2022).

8. P. K. Eide, V. Vinje, A. H. Pripp, K.-A. Mardal, G. Ringstad, Sleep deprivation impairs molecular clearance from the human brain. Brain 144, 863–874 (2021).

9. N. L. Hauglund, et al., Norepinephrine-mediated slow vasomotion drives glymphatic clearance during sleep. Cell 188, 606–622.e17 (2025).

10. T. Warbrick, Simultaneous EEG-fMRI: What Have We Learned and What Does the Future Hold? Sensors 22, 2262 (2022).

11. N. E. Fultz, et al., Coupled electrophysiological, hemodynamic, and cerebrospinal fluid oscillations in human sleep. Science (1979) 366, 628–631 (2019).

12. V. Kiviniemi, et al., Ultra-fast magnetic resonance encephalography of physiological brain activity – Glymphatic pulsation mechanisms? Journal of Cerebral Blood Flow & Metabolism 36, 1033–1045 (2016).

13. H. Helakari, et al., Human NREM Sleep Promotes Brain-Wide Vasomotor and Respiratory Pulsations. The Journal of Neuroscience 42, 2503–2515 (2022).

14. B. Battal, et al., Cerebrospinal fluid flow imaging by using phase-contrast MR technique. Br J Radiol 84, 758–765 (2011).

15. A. M. Wright, Y. Wu, L. Feng, Q. Wen, Diffusion magnetic resonance imaging of cerebrospinal fluid dynamics: Current techniques and future advancements. NMR Biomed 37 (2024).

16. M. de Zambotti, et al., K-Complexes: Interaction between the Central and Autonomic Nervous Systems during Sleep. Sleep 39, 1129–1137 (2016).

17. J. Tank, et al., Relationship between blood pressure, sleep K-complexes, and muscle sympathetic nerve activity in humans. *American Journal of Physiology-Regulatory*, Integrative and Comparative Physiology 285, R208–R214 (2003).

18. A. Lüthi, M. Nedergaard, Anything but small: Microarousals stand at the crossroad between noradrenaline signaling and key sleep functions. Neuron 113, 509–523 (2025).

19. A. Rechtschaffen, A. Kales, A manual of standardized terminology, techniques and scoring system of sleep stages in human subjects (Public Health Service, US Government Printing Office, 1968).

20. M. Boselli, L. Parrino, A. Smerieri, M. G. Terzano, Effect of Age on EEG Arousals in Normal Sleep. Sleep 21 (1998).

21. M. Uji, M. Tamaki, Sleep, learning, and memory in human research using noninvasive neuroimaging techniques. Neurosci Res 189, 66–74 (2023).

22. M. Tamaki, T. Watanabe, Y. Sasaki, Coregistration of magnetic resonance spectroscopy and polysomnography for sleep analysis in human subjects. STAR Protoc 2, 100974 (2021).

23. M. Tamaki, et al., Complementary contributions of non-REM and REM sleep to visual learning. Nat Neurosci 23, 1150–1156 (2020).

24. M. Tamaki, et al., First-night effect reduces the beneficial effects of sleep on visual plasticity and modifies the underlying neurochemical processes. Sci Rep 14, 14388 (2024).

25. X. Hu, T. H. Le, T. Parrish, P. Erhard, Retrospective estimation and correction of physiological fluctuation in functional MRI. Magn Reson Med 34, 201–212 (1995).

26. G. H. Glover, T.-Q. Li, D. Ress, Image-based method for retrospective correction of physiological motion effects in fMRI: RETROICOR. Magn Reson Med 44, 162–167 (2000).

27. O. C. Reddy, Y. D. van der Werf, The Sleeping Brain: Harnessing the Power of the Glymphatic System through Lifestyle Choices. Brain Sci 10, 868 (2020).

28. T. O. Wichmann, H. H. Damkier, M. Pedersen, A Brief Overview of the Cerebrospinal Fluid System and Its Implications for Brain and Spinal Cord Diseases. Front Hum Neurosci 15 (2022).

29. R. Berry, S. Quan, A. Abreu, Manual for the Scoring of Sleep and Associated Events: Rules, Terminology and Technical Specifications. American Academy of Sleep Medicine Version 2.6 (2020).

30. M. Massimini, R. Huber, F. Ferrarelli, S. Hill, G. Tononi, The Sleep Slow Oscillation as a Traveling Wave. The Journal of Neuroscience 24, 6862–6870 (2004).

31. T. T. Dang-Vu, et al., Spontaneous neural activity during human slow wave sleep. Proceedings of the National Academy of Sciences 105, 15160–15165 (2008).

32. M. Mölle, T. O. Bergmann, L. Marshall, J. Born, Fast and Slow Spindles during the Sleep Slow Oscillation: Disparate Coalescence and Engagement in Memory Processing. Sleep 34, 1411–1421 (2011).

33. V. Latreille, et al., The human K-complex: Insights from combined scalp-intracranial EEG recordings. Neuroimage 213, 116748 (2020).

34. Y. L. Wang, et al., Intracerebral dynamics of sleep arousals: a combined scalp-intracranial EEG study. The Journal of Neuroscience e0617232024 (2024). 10.1523/JNEUROSCI.0617-23.2024.

35. A. R. Adamantidis, C. Gutierrez Herrera, T. C. Gent, Oscillating circuitries in the sleeping brain. Nat Rev Neurosci 20, 746–762 (2019).

36. R. Boyce, S. D. Glasgow, S. Williams, A. Adamantidis, Causal evidence for the role of REM sleep theta rhythm in contextual memory consolidation. Science (1979) 352, 812– 816 (2016).

37. M. Dresler, et al., Neural Correlates of Dream Lucidity Obtained from Contrasting Lucid versus Non-Lucid REM Sleep: A Combined EEG/fMRI Case Study. Sleep 35, 1017–1020 (2012).

38. A. Elabasy, et al., Sleep increases propagation speed of physiological brain pulsations. [Preprint] (2025).

39. D. Picchioni, et al., Autonomic arousals contribute to brain fluid pulsations during sleep. Neuroimage 249, 118888 (2022).

40. S. D. Williams, et al., Neural activity induced by sensory stimulation can drive large-scale cerebrospinal fluid flow during wakefulness in humans. PLoS Biol 21, e3002035 (2023).

41. C. Strik, U. Klose, M. Erb, H. Strik, W. Grodd, Intracranial oscillations of cerebrospinal fluid and blood flows: Analysis with magnetic resonance imaging. Journal of Magnetic Resonance Imaging 15, 251–258 (2002).

42. L. Raitamaa, et al., Spectral analysis of physiological brain pulsations affecting the BOLD signal. Hum Brain Mapp 42, 4298–4313 (2021).

43. J. Gonzalez-Castillo, I. S. Fernandez, D. A. Handwerker, P. A. Bandettini, Ultra-slow fMRI fluctuations in the fourth ventricle as a marker of drowsiness. Neuroimage 259, 119424 (2022).

44. A. A. Borbely, A two process model of sleep regulation. Hum Neurobiol 1 (1982).

45. S. Brodt, M. Inostroza, N. Niethard, J. Born, Sleep—A brain-state serving systems memory consolidation. Neuron 111, 1050–1075 (2023).

46. G. Tononi, C. Cirelli, Sleep and the Price of Plasticity: From Synaptic and Cellular Homeostasis to Memory Consolidation and Integration. Neuron 81, 12–34 (2014).

47. M. Betta, et al., Cortical and subcortical hemodynamic changes during sleep slow waves in human light sleep. Neuroimage 236, 118117 (2021).

48. M. Schabus, et al., Hemodynamic cerebral correlates of sleep spindles during human non-rapid eye movement sleep. Proceedings of the National Academy of Sciences 104, 13164–13169 (2007).

49. D. Baena, et al., Functional differences in cerebral activation between slow wave-coupled and uncoupled sleep spindles. Front Neurosci 16 (2023).

50. R. Wehrle, et al., Functional microstates within human REM sleep: first evidence from fMRI of a thalamocortical network specific for phasic REM periods. European Journal of Neuroscience 25, 863–871 (2007).

51. T. T. Dang-Vu, et al., Functional Neuroimaging Insights into the Physiology of Human Sleep. Sleep 33, 1589–1603 (2010).

52. S. Miyauchi, M. Misaki, S. Kan, T. Fukunaga, T. Koike, Human brain activity time-locked to rapid eye movements during REM sleep. Exp Brain Res 192, 657–667 (2009).

53. S. Fogel, et al., Reactivation or transformation? Motor memory consolidation associated with cerebral activation time-locked to sleep spindles. PLoS One 12, e0174755 (2017).

54. M. Tamaki, T. Matsuoka, H. Nittono, T. Hori, Fast Sleep Spindle (13–15 Hz) Activity Correlates with Sleep-Dependent Improvement in Visuomotor Performance. Sleep 31, 204–211 (2008).

55. M. Schabus, et al., Interindividual sleep spindle differences and their relation to learning-related enhancements. Brain Res 1191, 127–135 (2008).

56. P. S. Özbay, et al., Sympathetic activity contributes to the fMRI signal. Commun Biol 2, 421 (2019).

57. L. Chen, A. Beckett, A. Verma, D. A. Feinberg, Dynamics of respiratory and cardiac CSF motion revealed with real-time simultaneous multi-slice EPI velocity phase contrast imaging. Neuroimage 122, 281–287 (2015).

58. V. Vijayakrishnan Nair, et al., Human CSF movement influenced by vascular low frequency oscillations and respiration. Front Physiol 13 (2022).

59. I. Slutsky, Linking activity dyshomeostasis and sleep disturbances in Alzheimer disease. Nat Rev Neurosci 25, 272–284 (2024).

60. Y.-E. S. Ju, et al., Sleep Quality and Preclinical Alzheimer Disease. JAMA Neurol 70, 587 (2013).

61. M. V. Vitiello, P. N. Prinz, D. E. Williams, M. S. Frommlet, R. K. Ries, Sleep Disturbances in Patients With Mild-Stage Alzheimer’s Disease. J Gerontol 45, M131–M138 (1990).

62. B. A. Mander, J. R. Winer, M. P. Walker, Sleep and Human Aging. Neuron 94, 19–36 (2017).

63. G. Liu, A. Ladrón-de-Guevara, Y. Izhiman, M. Nedergaard, T. Du, Measurements of cerebrospinal fluid production: a review of the limitations and advantages of current methodologies. Fluids Barriers CNS 19, 101 (2022).

64. L. Bojarskaite, et al., Sleep cycle-dependent vascular dynamics in male mice and the predicted effects on perivascular cerebrospinal fluid flow and solute transport. Nat Commun 14, 953 (2023).

65. L.-F. Jiang-Xie, et al., Neuronal dynamics direct cerebrospinal fluid perfusion and brain

66. T. C. Diorio, et al., Real-time quantification of in vivo cerebrospinal fluid velocity using the functional magnetic resonance imaging inflow effect. NMR Biomed 37 (2024).

67. V. V. Nair, T. C. Diorio, Q. Wen, V. L. Rayz, Y. Tong, Using respiratory challenges to modulate CSF movement across different physiological pathways: An fMRI study. Imaging Neuroscience 2, 1–14 (2024).

68. T. Martins, et al., Characterization of pulsations in the brain and cerebrospinal fluid using ultra-high field magnetic resonance imaging. Front Neurosci 18 (2024).

69. J. Tuunanen, et al., Cardiovascular and vasomotor pulsations in the brain and periphery during awake and NREM sleep in a multimodal fMRI study. Front Neurosci 18 (2024).

70. H. Helakari, et al., Effect of sleep deprivation and NREM sleep stage on physiological brain pulsations. Front Neurosci 17 (2023).

71. C. Song, M. Boly, E. Tagliazucchi, H. Laufs, G. Tononi, fMRI spectral signatures of sleep. Proceedings of the National Academy of Sciences 119 (2022).

72. D. J. Buysse, C. F. Reynolds, T. H. Monk, S. R. Berman, D. J. Kupfer, The Pittsburgh sleep quality index: A new instrument for psychiatric practice and research. Psychiatry Res 28, 193–213 (1989).

73. E. Hoddes, V. Zarcone, H. Smythe, R. Phillips, W. C. Dement, Quantification of Sleepiness: A New Approach. Psychophysiology 10, 431–436 (1973).

74. M. Uji, R. Wilson, S. T. Francis, K. J. Mullinger, S. D. Mayhew, Exploring the advantages of multiband fMRI with simultaneous EEG to investigate coupling between gamma frequency neural activity and the BOLD response in humans. Hum Brain Mapp 39, 1673– 1687 (2018).

75. P. J. Allen, O. Josephs, R. Turner, A Method for Removing Imaging Artifact from Continuous EEG Recorded during Functional MRI. Neuroimage 12, 230–239 (2000).

76. R. Wilson, K. J. Mullinger, S. T. Francis, S. D. Mayhew, The relationship between negative BOLD responses and ERS and ERD of alpha/beta oscillations in visual and motor cortex. Neuroimage 199, 635–650 (2019).

77. M. Bullock, G. D. Jackson, D. F. Abbott, Artifact Reduction in Simultaneous EEG-fMRI: A Systematic Review of Methods and Contemporary Usage. Front Neurol 12 (2021).

78. R. Agarwal, T. Takeuchi, S. Laroche, J. Gotman, Detection of Rapid-Eye Movements in Sleep Studies. IEEE Trans Biomed Eng 52, 1390–1396 (2005).

79. B. Frauscher, et al., Rapid Eye Movement Sleep Sawtooth Waves Are Associated with Widespread Cortical Activations. The Journal of Neuroscience 40, 8900–8912 (2020).

80. L. Peter-Derex, et al., Enhanced thalamocortical functional connectivity during rapid-eye-movement sleep sawtooth waves. Sleep 46 (2023).

81. S. Sato, et al., Relationship between muscle tone changes, sawtooth waves and rapid eye movements during sleep. Electroencephalogr Clin Neurophysiol 103, 627–632 (1997).

82. B. Fischl, FreeSurfer. Neuroimage 62, 774–781 (2012).

83. B. Fischl, A. M. Dale, Measuring the thickness of the human cerebral cortex from magnetic resonance images. Proceedings of the National Academy of Sciences 97, 11050–11055 (2000).

84. M. Jenkinson, P. Bannister, M. Brady, S. Smith, Improved Optimization for the Robust and Accurate Linear Registration and Motion Correction of Brain Images. Neuroimage 17, 825–841 (2002).

85. S. M. Smith, Fast robust automated brain extraction. Hum Brain Mapp 17, 143–155 (2002).

86. R. Scheeringa, et al., Neuronal Dynamics Underlying High– and Low-Frequency EEG Oscillations Contribute Independently to the Human BOLD Signal. Neuron 69, 572–583 (2011).

87. R. Oostenveld, P. Fries, E. Maris, J.-M. Schoffelen, FieldTrip: Open Source Software for Advanced Analysis of MEG, EEG, and Invasive Electrophysiological Data. Comput Intell Neurosci 2011, 1–9 (2011).

88. T. Donoghue, et al., Parameterizing neural power spectra into periodic and aperiodic

89. M. Fukunaga, et al., Large-amplitude, spatially correlated fluctuations in BOLD fMRI signals during extended rest and early sleep stages. Magn Reson Imaging 24, 979–992 (2006).

90. S. G. Horovitz, et al., Low frequency BOLD fluctuations during resting wakefulness and light sleep: A simultaneous EEG-fMRI study. Hum Brain Mapp 29, 671–682 (2008).

91. S. Olbrich, et al., EEG-vigilance and BOLD effect during simultaneous EEG/fMRI measurement. Neuroimage 45, 319–332 (2009).

92. L. J. Larson-Prior, et al., Cortical network functional connectivity in the descent to sleep. Proceedings of the National Academy of Sciences 106, 4489–4494 (2009).

93. C. Caballero-Gaudes, R. C. Reynolds, Methods for cleaning the BOLD fMRI signal. Neuroimage 154, 128–149 (2017).

94. M. W. Woolrich, T. E. J. Behrens, C. F. Beckmann, M. Jenkinson, S. M. Smith, Multilevel linear modelling for FMRI group analysis using Bayesian inference. Neuroimage 21, 1732–1747 (2004).

95. E. T. Rolls, C.-C. Huang, C.-P. Lin, J. Feng, M. Joliot, Automated anatomical labelling atlas 3. Neuroimage 206, 116189 (2020).

96. E. Maris, Statistical testing in electrophysiological studies. Psychophysiology 49, 549–565 (2012).

97. E. Maris, R. Oostenveld, Nonparametric statistical testing of EEG– and MEG-data. J Neurosci Methods 164, 177–190 (2007).

98. T. E. Nichols, A. P. Holmes, Nonparametric permutation tests for functional neuroimaging: A primer with examples. Hum Brain Mapp 15, 1–25 (2002).

99. C. R. Genovese, N. A. Lazar, T. Nichols, Thresholding of Statistical Maps in Functional Neuroimaging Using the False Discovery Rate. Neuroimage 15, 870–878 (2002).

