## Supplemental information for "Human deep sleep facilitates faster cerebrospinal fluid dynamics linked to brain oscillations for sleep homeostasis and memory"

**This PDF file includes:**

Figures S1 to S12
Tables S1 to S9

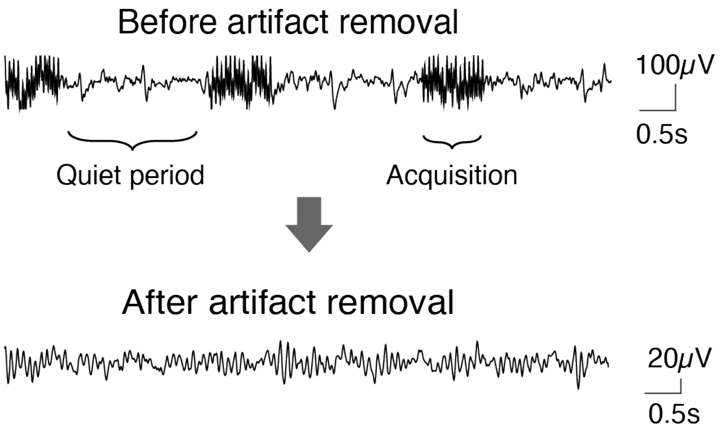


**Fig. S1.** Example EEG data before and after artifact removal obtained via the sparse fMRI method developed in the present study. See **MRI acquisition, PSG measurement,** and **PSG data preprocessing** sections in Materials and Methods for details.

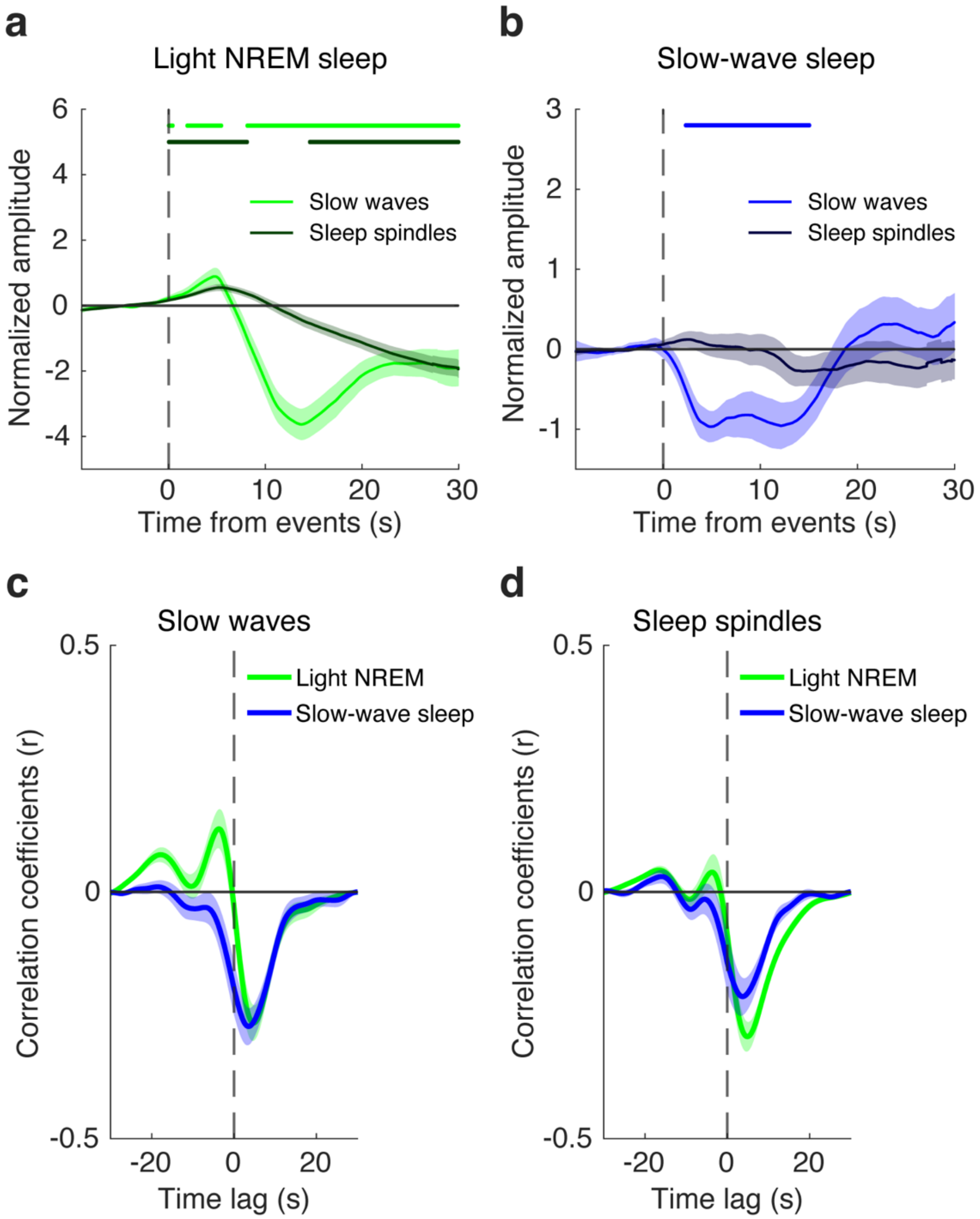


**Fig. S2. GM signal changes time-locked to slow waves and sleep spindles. a**, GM signal amplitude changes time-locked to slow waves (light green) and sleep spindles (dark green) during light NREM sleep. The horizontal bars (light green: slow waves; dark green, sleep spindles) indicate significances against zero baselines using two-tailed one-sample t-tests, (FDR corrected, *Ps* < 0.05). **b**, GM signal amplitude changes time-locked to slow waves (light blue) and sleep spindles (dark blue) during slow-wave sleep. The horizontal bar (light blue, slow waves) indicates significances against zero baselines using two-tailed one-sample t-tests, (FDR corrected, *Ps* < 0.05). **c**, Cross-correlation between CSF and GM signals time-locked to slow waves (light NREM: max |r| = -0.27, at lag = 4.8s; SWS: max |r| = -0.27, at lag = 3.5s). Green, light NREM sleep. Blue, slow-waves sleep. **d**, Cross-correlation between CSF and GM signals time-locked to sleep spindles (light NREM: max |r| = -0.29, at lag = 4.9s; SWS: max |r| = -0.21, at lag = 3.7s). Green, light NREM sleep. Blue, slow-waves sleep.

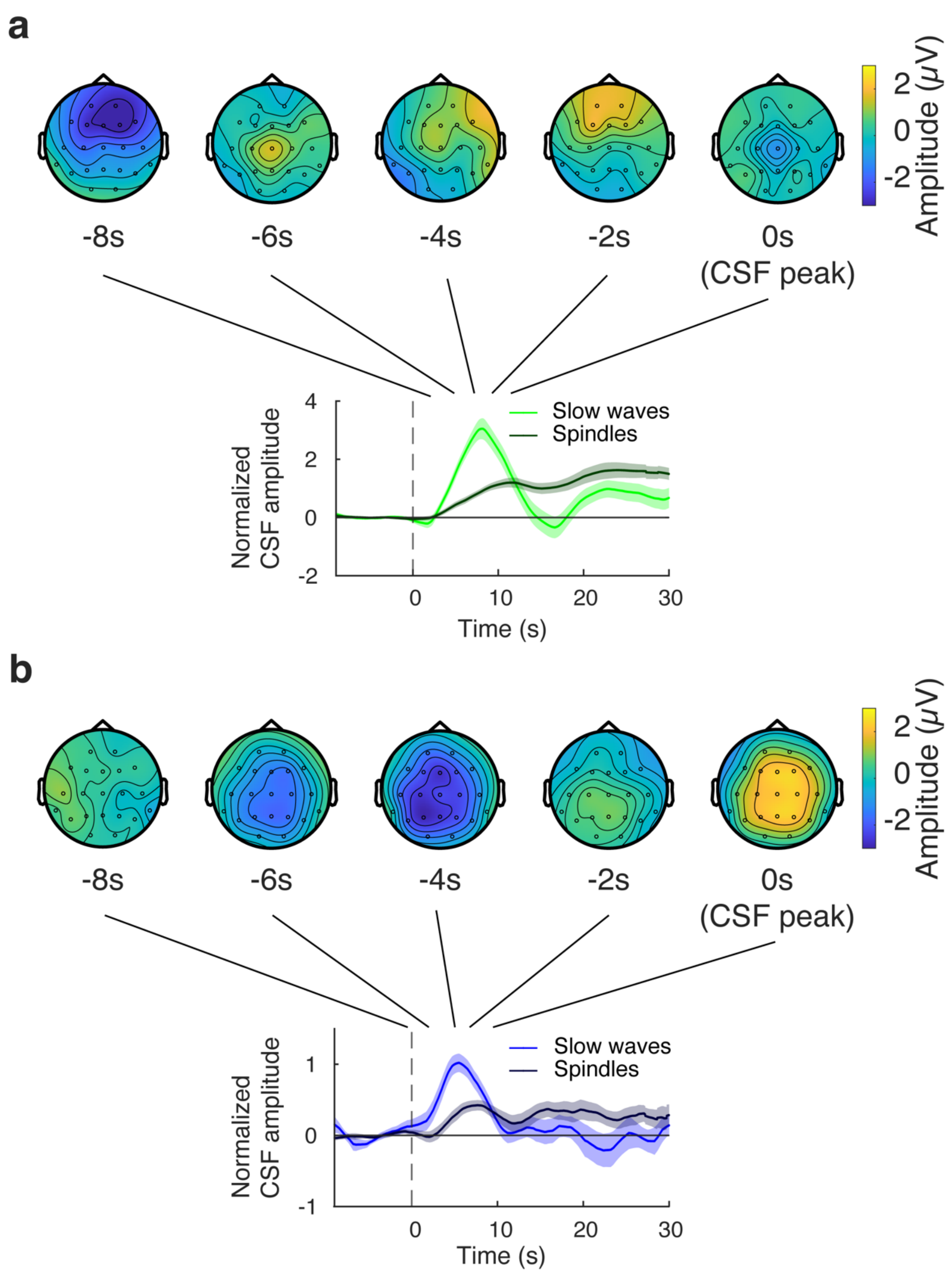


**Fig. S3. EEG topographical changes time-locked to CSF signal peaks during light NREM (a) and slow-wave sleep (b).** t=0s corresponds to the CSF signal peak. **(a)** EEG topographic map indicated a negative peak centered around the frontal region, resembling a topography of K-complexes, which peaked 8 s *before* the CSF positive peak events during light NREM sleep. **(b)** A negative peak spread through the midline of frontal to parietal regions resembling those found typically in slow waves (1) which peaked around 5 s *before* the CSF positive peak events during slow-wave sleep. Note that slow waves during slow-wave sleep were followed by a CSF peak 5.5s *after* the onset of the events. The plots for CSF signals are the same as those in **Fig. 1d** and **e**.

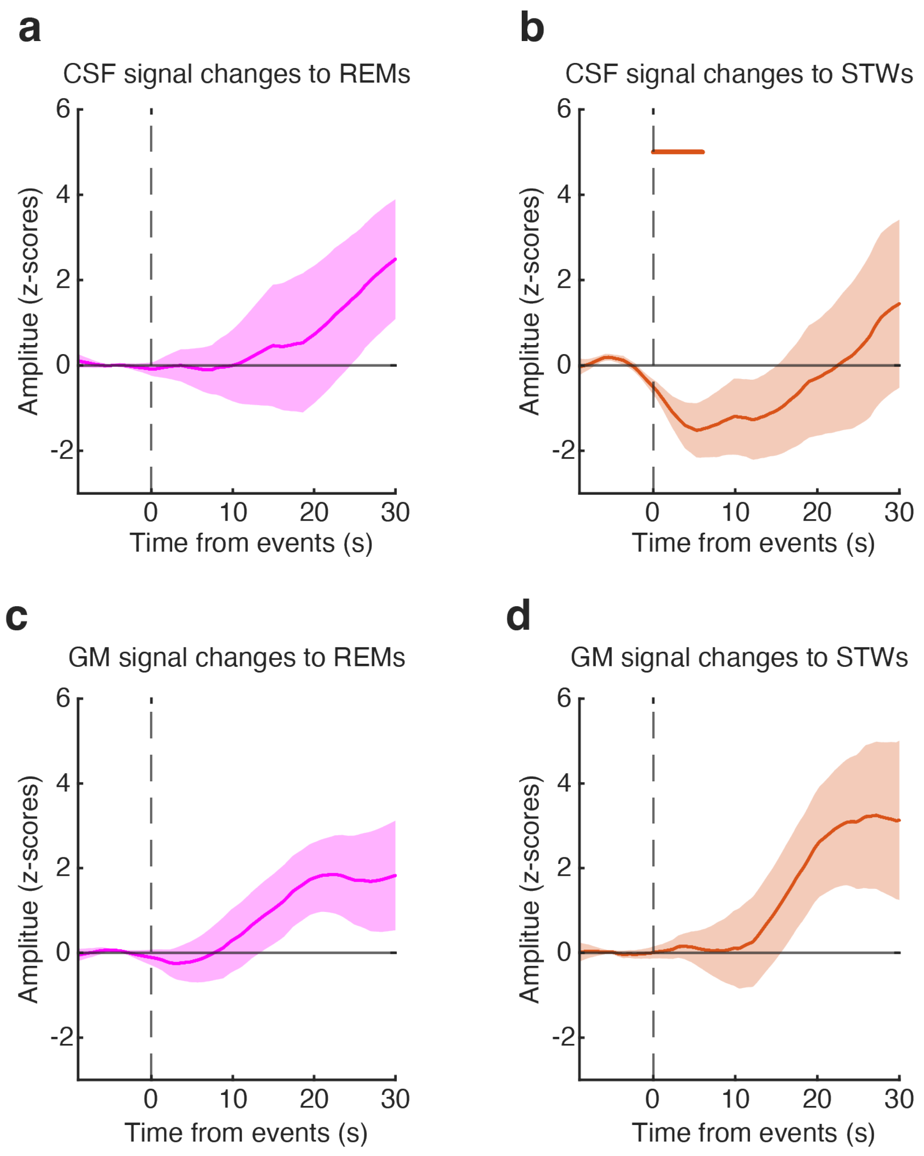


**Fig. S4. CSF and GM signal changes time-locked to events during REM sleep. a**, CSF signal changes to rapid eye movements (REMs). **b**, CSF signal changes to sawtooth waves (STWs). The horizontal bar (orange, sawtooth waves) indicates significances against zero baselines using two-tailed one-sample t-tests, (FDR corrected, *Ps* < 0.05). **c**, GM signal changes to rapid eye movements (REMs). **d**, GM signal changes to sawtooth waves (STWs).

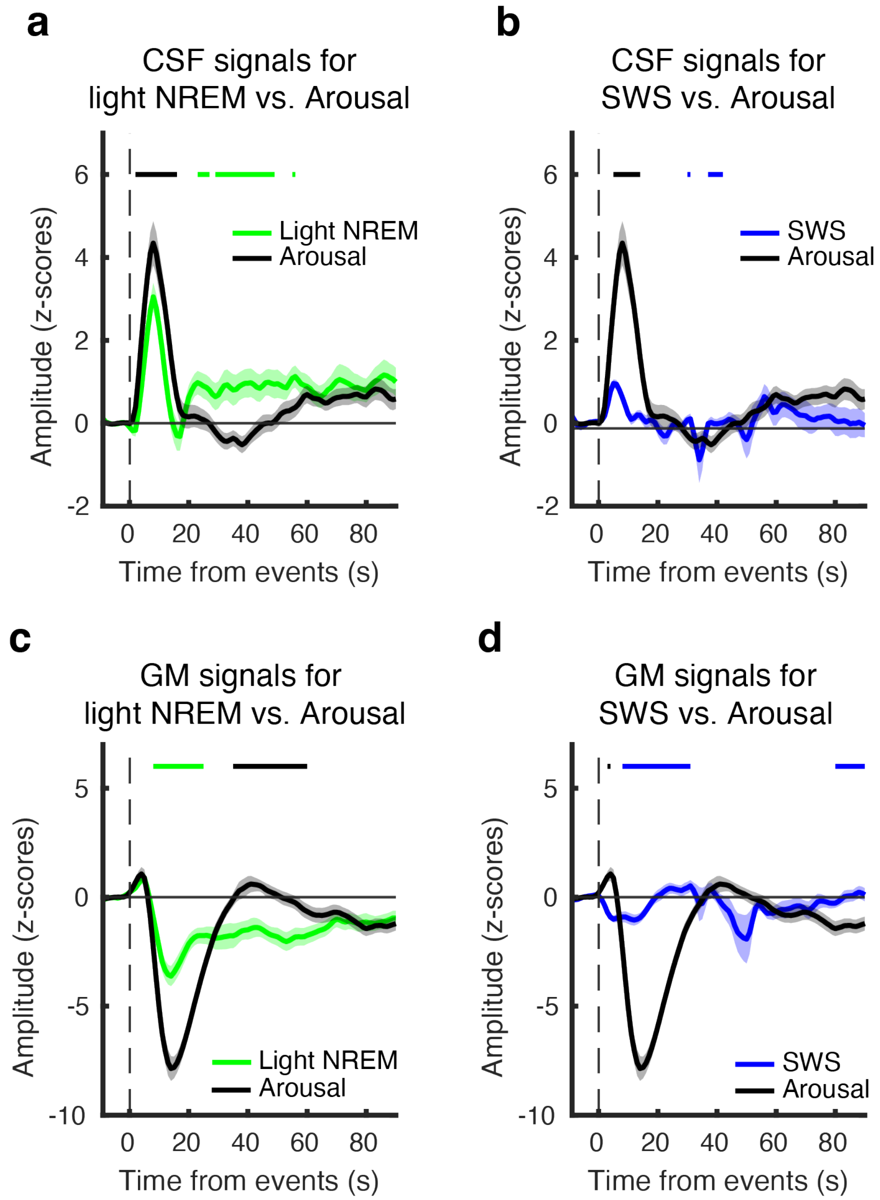


**Fig. S5. Comparisons of CSF and GM signal changes following slow waves in different sleep depths and arousals. a & b**, CSF signal changes time-locked to slow waves during sleep and arousals. Green, light NREM sleep; blue, slow-wave sleep; black, arousals. **c & d**, GM signal changes time-locked to slow waves during sleep and arousals. Green, light NREM sleep, blue, slow-wave sleep, black, arousals. The horizontal bars (green: light NREM; blue, slow-wave sleep; black: arousals) indicate significant differences between possible pairs using two-tailed paired *t*-tests, (FDR corrected, *Ps* < 0.05). SWS, slow-wave sleep.

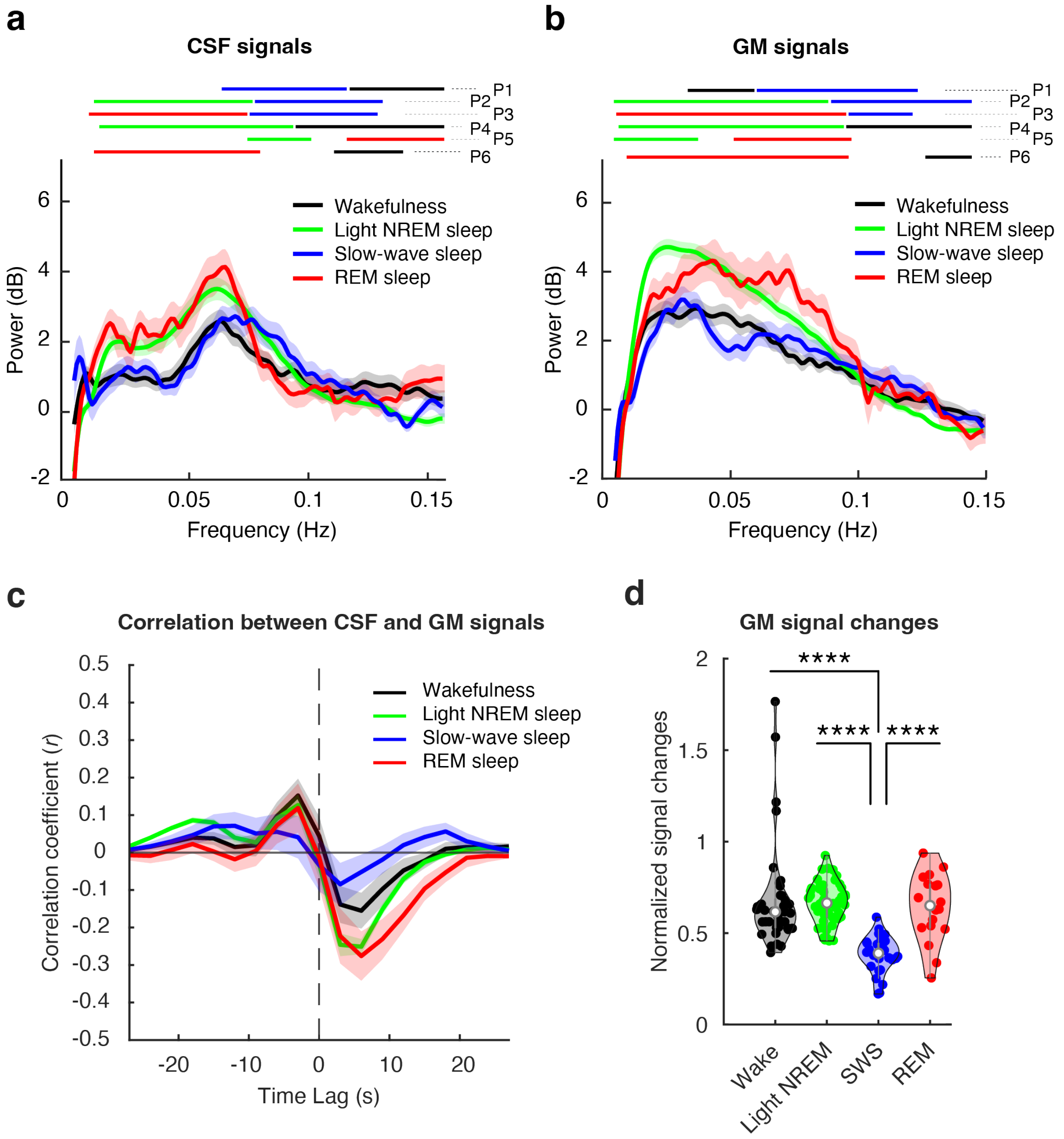


**Fig. S6. Frequency profiles of CSF signals in the lateral ventricles and GM signals vary across different sleep depths. a,** Periodic CSF signal power spectrum against each frequency bin for each sleep stage. Each of the horizontal bars at the top represent a significant cluster for possible pairs (P1~6) where the color indicates the significantly larger power in the corresponding stage compared to the other stage pair (cluster-based permutation tests, FDR-corrected *Ps* < 0.05). The faster component power at approximately 0.06-0.12Hz was significantly greater during slow-wave sleep than during wakefulness (P1, blue vs. black; FDR-corrected $P < 0.05$) or light NREM sleep (P2, blue vs. green; FDR-corrected $P < 0.05$). In contrast, the slower component power (approximately 0.06 Hz or below) was significantly larger during light NREM than during wakefulness (P4, green vs. black; FDR-corrected $P < 0.05$) or slow-wave sleep (P2, green vs. blue; FDR-corrected $P < 0.05$). During REM sleep, slower components were significantly larger than during slow-wave sleep and wakefulness (P3, red vs. blue; P6, red vs. black; FDR-corrected *Ps* < 0.05) and very fast component (above 0.12 Hz) were significantly larger than during light NREM sleep (P5, red vs. green; FDR-corrected $P < 0.05$). Pair (P) 1, Slow-wave sleep (blue) vs. wakefulness (black). P2, Slow-wave sleep vs. light NREM sleep (green). P3, Slow-wave sleep vs. REM sleep. P4, Light NREM sleep vs. wakefulness. P5, Light NREM sleep vs. REM sleep. P6, REM sleep vs. wakefulness. **b,** Periodic GM signal power spectrum against each frequency bin for each sleep stage. **c**, Cross-correlation between CSF and GM signals (Wake: max r = 0.15 at lag = -3s, min r = -0.16 at lag = 6s; light NREM: max r = 0.12 at lag = -3s, min r = -0.25 at lag = 3s; SWS: max r = 0.07 at lag = -12s, min r = -0.08 at lag = 3s; REM: max r = 0.12 at lag = -3s, min r = -0.28 at lag = 6s). **d,** GM signal variances during different sleep stages. Two-tailed paired *t*-test with Bonferroni correction, ****$P < 0.001$

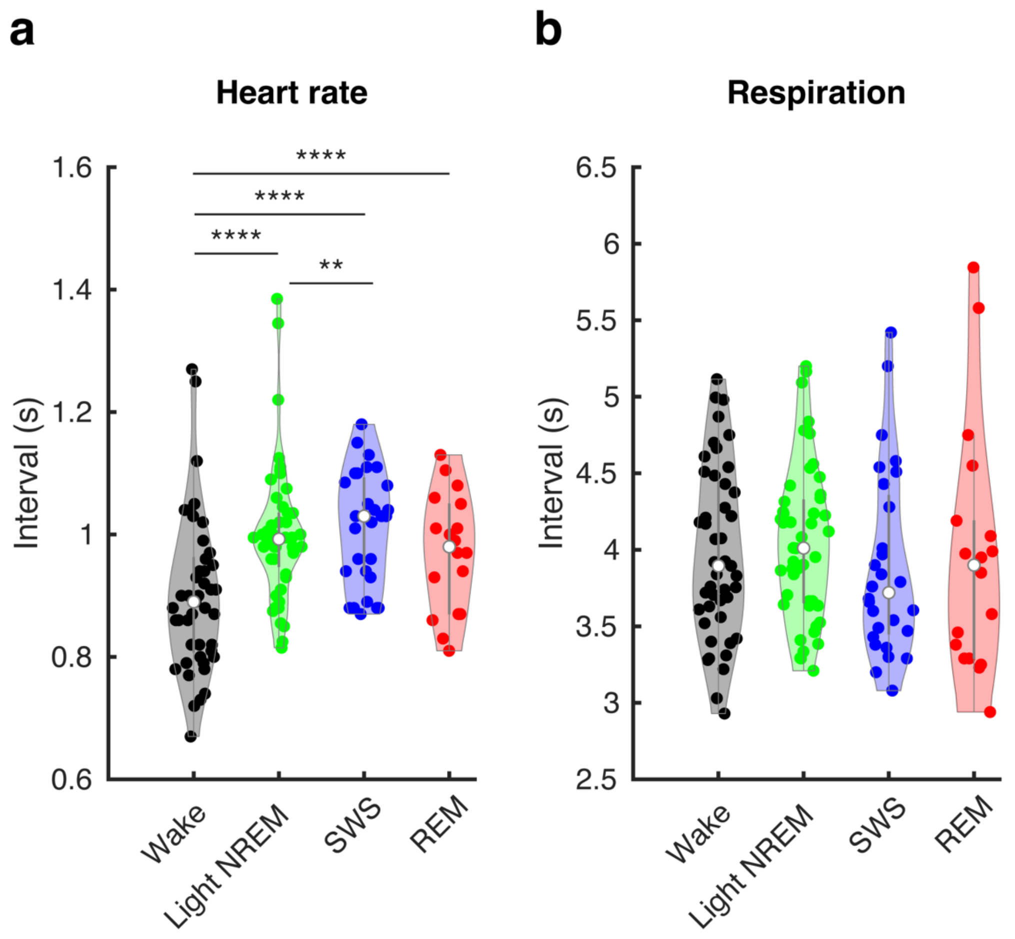


**Fig. S7. Intervals of cardiac and respiratory cycles during sleep and wakefulness. a**, Intervals of cardiac cycles. The intervals of the cardiac cycles were significantly longer during sleep than during wakefulness (light NREM sleep vs. wakefulness: *P* = 5.1303e-09; Slow-wave sleep vs. wakefulness: *P* = 1.2157e-05; REM sleep vs. wakefulness: *P* = 9.7656e-04). Slow-wave sleep had significantly longer intervals than light NREM (*P* = 0.0016). Two-tailed Wilcoxson signed-rank test with Bonferroni correction, ****$P$<.001, **$P$<.01. **b**, Intervals of respiratory cycles. There was no significant difference in the intervals of respiratory cycles across sleep stages. SWS, slow-wave sleep.

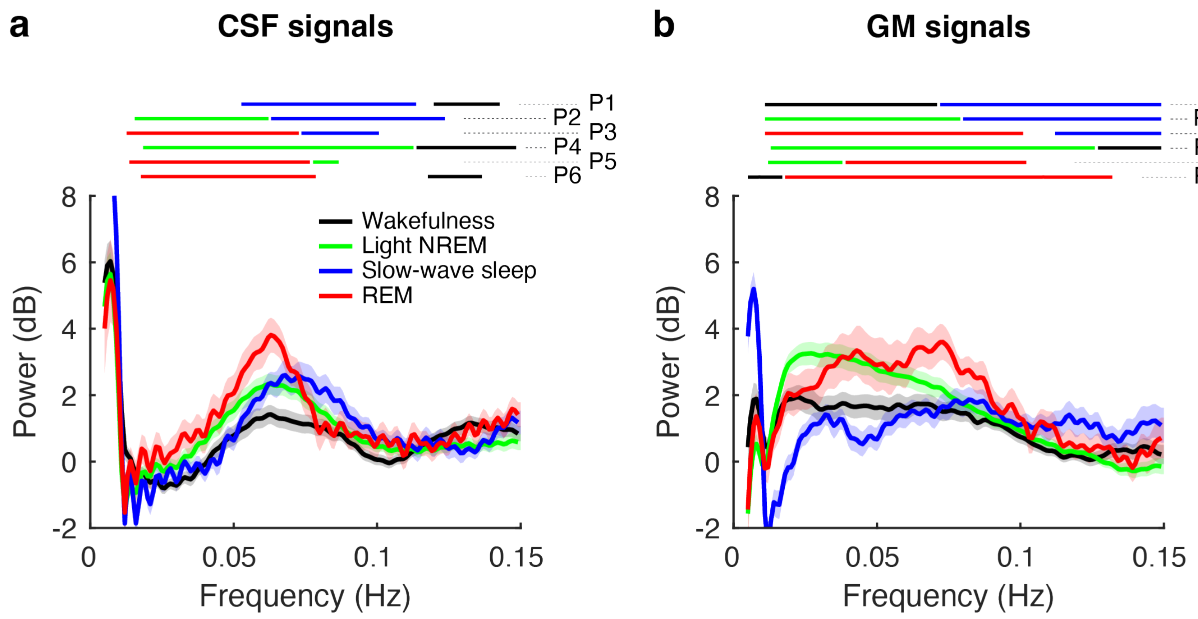


**Fig. S8. Periodic frequency profiles obtained from the CSF (a) and GM (b) signals without denoising cardiac and respiration signal contaminations.** Periodic CSF signal power spectrum against each frequency bin for each sleep stage. Each of the horizontal bars at the top represent a significant cluster for possible pairs (P1~6) where the color indicates the significantly larger power in the corresponding stage compared to the other stage pair (cluster-based permutation tests, *Ps* < 0.05). Pair (P) 1, Slow-wave sleep (blue) vs. wakefulness (black). P2, Slow-wave sleep vs. light NREM sleep (green). P3, Slow-wave sleep vs. REM sleep. P4, Light NREM sleep vs. wakefulness. P5, Light NREM sleep vs. REM sleep. P6, REM sleep vs. wakefulness.

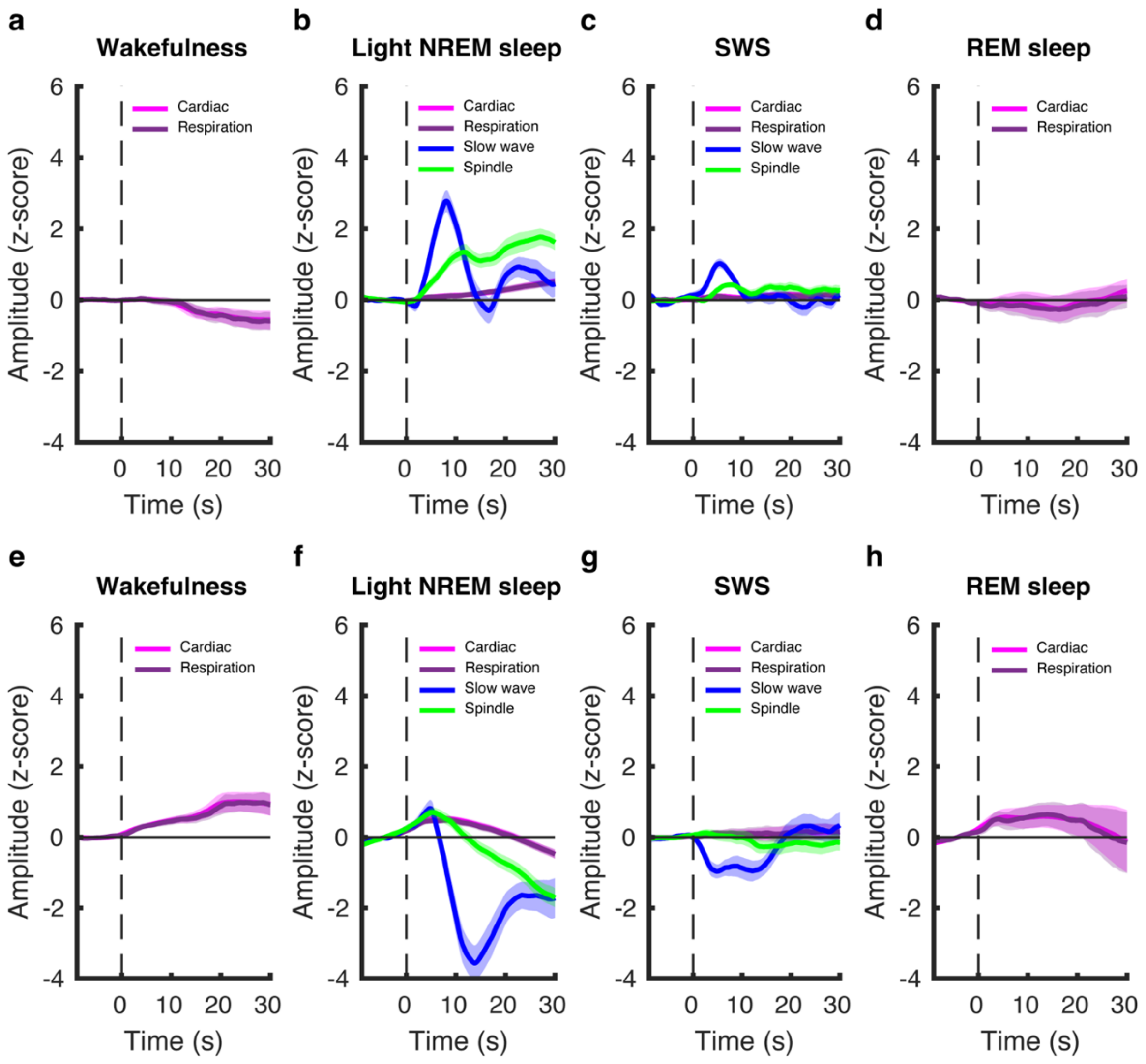


**Fig. S9. Comparisons of CSF and GM signal changes induced by cardiac pulses and respirations vs. slow waves and sleep spindles in different sleep depths.** We investigated the CSF signal changes time-locked to respiration and cardiac pulses in the same manner as in previous studies (2, 3). **a–d**, CSF signal changes time-locked to each event onset during each sleep stage. **e–h**, GM signal changes time-locked to each event onset during each sleep stage. Magenta, cardiac pulses. Purple, respirations, Blue, slow waves, Green, spindles. SWS, slow-wave sleep.

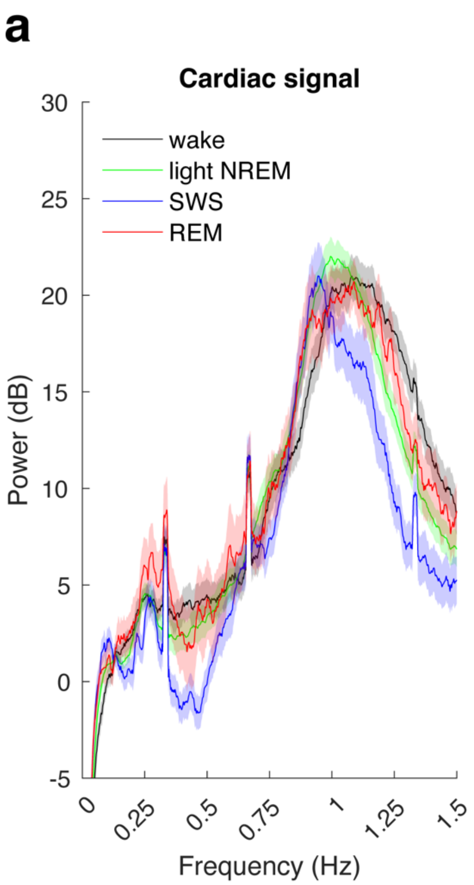


**Fig. S10. Periodic frequency profiles in the cardiac cycles (a) and the respiration cycles (b).** The power spectrum of the cardiac and respiration signals was measured using the same analysis steps as the power spectrum analysis of fMRI data. There was a peak power at around 1 Hz in the cardiac signals, whereas a peak power of around 0.3 Hz was observed in the respiration signals in accordance with a previous study (4). Thus, the cardiac and respirations had much higher frequency signals (>0.3Hz) that did not overlap with the frequency range induced by sleep brain oscillations (0.06–0.15Hz, **Fig. 1d, e**).

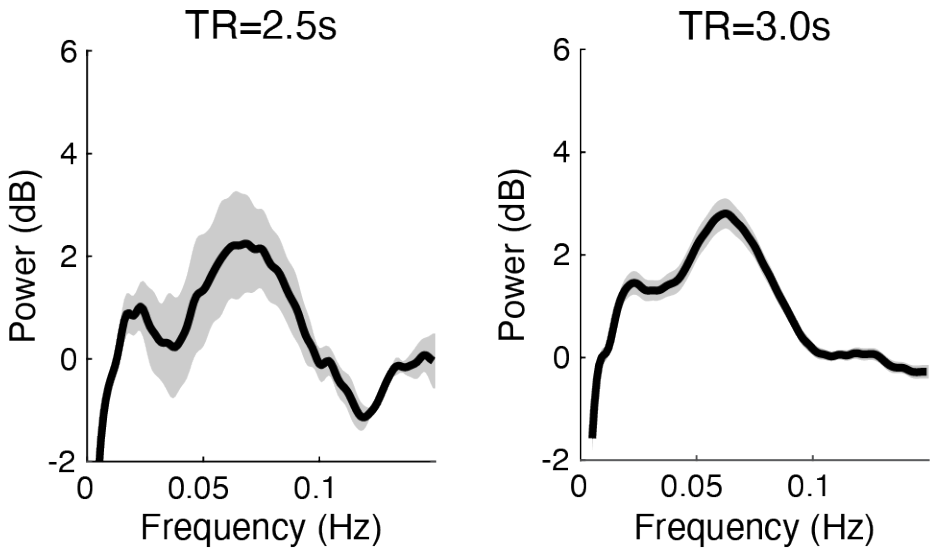


**Fig. S11.** Group mean power spectrum of CSF signals measured by TR=2.5s (n=3 nap sessions, left) and TR=3.0s (n=50 nap sessions, right). See ***The control experiment and analyses to test aliasing noises*** in **Materials and Methods** for more details.

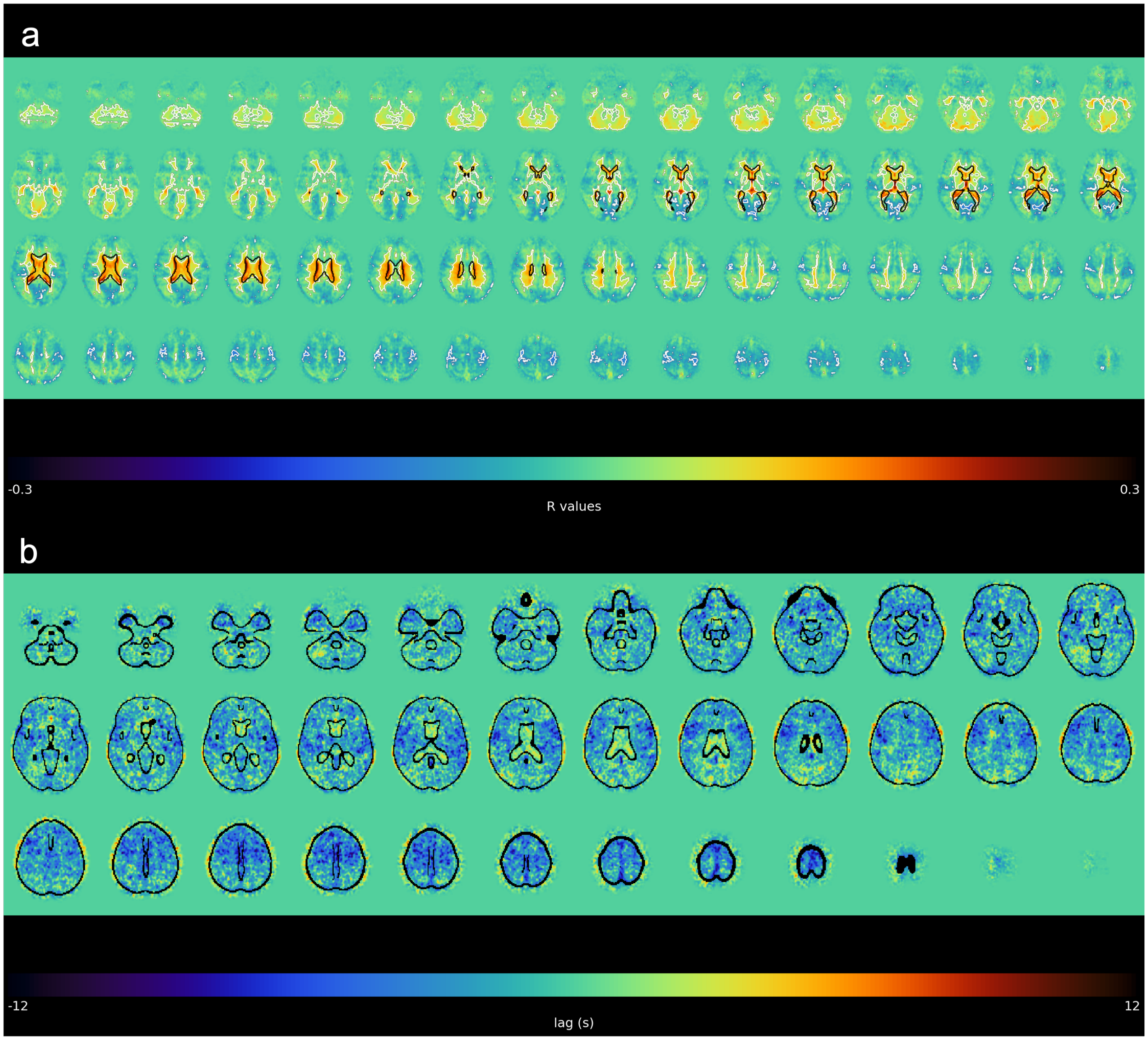


**Fig. S12. Relationship between the lateral ventricle signals and fMRI signals in other regions. a,** Group mean whole brain voxel-wise zero-lag correlation map between the CSF signals from lateral ventricles and fMRI signals in each voxel. Correlation coefficient values were calculated by Pearson's correlation with the seed signal, which was the average signal of the lateral ventricles. We found positive correlations within the ventricular regions, including the lateral and the fourth ventricles (warmer colors). We also observed wide-spread negative correlations between the lateral ventricle signals and grey matter regions (colder colors). Importantly, the strongest positive correlations ($r = 0.3$) concentrate in the lateral ventricles. Black outlines highlight lateral ventricle regions. White outlines highlight significant voxels against zero correlation coefficient values (uncorrected $p<0.001$). **b,** Group mean whole brain voxel-wise time-lag map of the maximum positive correlation between the CSF signals from lateral ventricles and fMRI signals in each voxel. Time-lags in each voxel were calculated by cross-correlations between the seed signal, which was the average signal of the lateral ventricles, and fMRI signals in each voxel for each nap session. Colder colors indicate that fMRI signals at that location precede the seed signal (i.e. lateral ventricle signal), whereas warmer colors indicate the lateral ventricle signal precedes fMRI signals at that location. We found that grey matter regions generally precede lateral ventricle signals (colder colors), whereas small lags can be found within in the ventricular regions. Additionally, positive lags (warmer colors) can be found in the subarachnoid space, meaning the lateral ventricle signals precede the subarchnoid changes. Black lines outline the structure of brain.

**Table S1. Sleep parameters**

| | NP | NS | Mean ± SE | | |
|---|---|---|---|---|---|
| Time in bed (min) | 25 | 50 | 89.7 | ± | 0.3 |
| Sleep efficiency (%) | 25 | 50 | 77.6 | ± | 3.2 |
| Wakefulness (%) | 25 | 48 | 22.4 | ± | 3.2 |
| Light NREM sleep (%) | 25 | 50 | 62.8 | ± | 2.9 |
| Slow-wave sleep (%) | 18 | 30 | 12.3 | ± | 2.3 |
| REM sleep (%) | 13 | 18 | 2.4 | ± | 0.6 |

**NP**, the number of participants. **NS**, the number of sessions. Time in bed indicates the duration of sleep sessions during MRI scan (the time interval between lights-off and lights-on). Sleep efficiency was measured by [total sleep duration / time in bed]. Light NREM sleep indicates the percentage of time in NREM sleep stages 1 and 2. Slow-wave sleep indicates the percentage of time in NREM sleep stage 3. REM sleep indicates the percentage of time in REM sleep stage. PSG data were scored in accordance with standard criteria (5).

**Table S2:** Summary of key oscillations, recruited brain regions, and CSF characteristics during sleep and arousals.

| | **Light NREM sleep** | **Slow-wave sleep** | **REM sleep** | **Arousals** |
|---|---|---|---|---|
| Key oscillations / events | K-complexes<br>Sleep spindles | Slow waves<br>Sleep spindles | Rapid eye movements<br>Sawtooth waves | N/A (mixed) |
| Brain regions | Sensory | Frontal, hippocampus | Visual, hippocampus | Widespread |
| CSF frequency | Slow | Fast | Slow<br>very fast | N/A |
| CSF amplitude | Larger | Smaller | Larger | Larger |
| CSF latency | Longer | Shorter | Longer | Longer |

Brain regions indicate activated brain regions to key oscillations or events in each sleep stage. CSF power frequency indicates the frequency characteristics in each sleep stage. CSF amplitude and latency indicates relative amplitudes and latencies in the CSF signals time-locked to the onset of brain oscillations / events across different sleep stages. **Green** cells indicate slower and larger impacts on the CSF signals, representing strong but infrequent CSF and brain dynamics. **Blue** cells indicate faster and smaller impacts on the CSF signals, representing mild but frequent CSF and brain dynamics. **Red** cell indicates a very fast component in the CSF signals that is specific to REM sleep with a mixture of the slow component (green).

**Table S3.** Brain regions that significantly increased for slow waves during light NREM sleep than during slow-wave sleep.

| ROIs | *Z* values | *P* values |
|---|---|---|
| Cingulate_Mid_L | 3.8082 | 0.0002 |
| Cingulate_Mid_R | 3.6531 | 0.0002 |
| Cingulate_Post_L | 3.6264 | 0.0002 |
| Cingulate_Post_R | 3.3095 | 0.0006 |
| Calcarine_L | 3.7446 | 0.0002 |
| Calcarine_R | 3.6056 | 0.0003 |
| Cuneus_L | 3.7074 | 0.0003 |
| Cuneus_R | 3.5805 | 0.0003 |
| Lingual_L | 3.7504 | 0.0002 |
| Lingual_R | 3.7962 | 0.0003 |
| Occipital_Sup_L | 3.5848 | 0.0003 |
| Occipital_Sup_R | 3.5587 | 0.0003 |
| Occipital_Mid_L | 3.8243 | 0.0002 |
| Occipital_Mid_R | 3.6286 | 0.0003 |
| Occipital_Inf_L | 3.8286 | 0.0002 |
| Occipital_Inf_R | 3.5069 | 0.0004 |
| Fusiform_L | 3.5723 | 0.0003 |
| Fusiform_R | 3.2599 | 0.0006 |
| Postcentral_L | 3.7671 | 0.0002 |
| Parietal_Sup_L | 3.6435 | 0.0003 |
| Parietal_Sup_R | 3.9973 | 0.0002 |
| Parietal_Inf_L | 3.4056 | 0.0004 |
| Parietal_Inf_R | 3.3055 | 0.0005 |
| Angular_L | 3.4215 | 0.0004 |
| Angular_R | 3.4098 | 0.0005 |
| Precuneus_L | 4.1504 | 0.0001 |
| Precuneus_R | 3.7021 | 0.0003 |
| Paracentral_Lobule_L | 3.3912 | 0.0005 |
| Temporal_Mid_L | 3.5801 | 0.0003 |
| Temporal_Mid_R | 3.7628 | 0.0002 |
| Temporal_Inf_L | 3.7744 | 0.0002 |
| Temporal_Inf_R | 3.7552 | 0.0002 |
| Cerebellum_Crus1_L | 3.6657 | 0.0003 |
| Cerebellum_Crus1_R | 3.1931 | 0.0007 |
| Cerebellum_4_5_L | 3.5464 | 0.0003 |
| Cerebellum_4_5_R | 3.3023 | 0.0006 |
| Cerebellum_6_L | 3.6276 | 0.0001 |
| Cerebellum_6_R | 3.4245 | 0.0004 |
| Vermis_4_5 | 3.5228 | 0.0003 |

Each ROI label is in accordance with the automated anatomical atlas 3 (6). L, left hemisphere. R, right hemisphere.

**Table S4.** Brain regions that significantly increased for slow waves during slow-wave sleep than during light NREM sleep.

| ROIs | *Z* values | *P* values |
|---|---|---|
| Precentral_L | 3.7206 | 0.0003 |
| Precentral_R | 3.7044 | 0.0003 |
| Frontal_Sup_2_L | 3.6866 | 0.0003 |
| Frontal_Sup_2_R | 3.9108 | 0.0002 |
| Frontal_Mid_2_L | 3.3904 | 0.0004 |
| Frontal_Mid_2_R | 3.3258 | 0.0005 |
| Frontal_Inf_Oper_L | 4.3669 | 0.0001 |
| Frontal_Inf_Oper_R | 4.0255 | 0.0002 |
| Frontal_Inf_Tri_L | 4.1318 | 0.0002 |
| Frontal_Inf_Tri_R | 3.8356 | 0.0002 |
| Frontal_Inf_Orb_2_L | 4.1623 | 0.0001 |
| Frontal_Inf_Orb_2_R | 4.2988 | 0.0002 |
| Rolandic_Oper_L | 3.5048 | 0.0003 |
| Rolandic_Oper_R | 3.6381 | 0.0003 |
| Supp_Motor_Area_L | 3.9727 | 0.0002 |
| Supp_Motor_Area_R | 3.9846 | 0.0002 |
| Olfactory_L | 3.4511 | 0.0004 |
| Olfactory_R | 3.7452 | 0.0003 |
| Frontal_Sup_Medial_L | 3.9463 | 0.0002 |
| Frontal_Sup_Medial_R | 4.0646 | 0.0002 |
| Frontal_Med_Orb_L | 3.9534 | 0.0002 |
| Frontal_Med_Orb_R | 4.3550 | 0.0001 |
| Rectus_L | 3.8079 | 0.0003 |
| Rectus_R | 3.5458 | 0.0003 |
| OFCmed_L | 3.4402 | 0.0005 |
| OFCmed_R | 3.2660 | 0.0006 |
| OFCant_L | 3.4433 | 0.0004 |
| OFCpost_L | 3.9637 | 0.0002 |
| OFCpost_R | 4.0323 | 0.0002 |
| OFClat_L | 3.2987 | 0.0005 |
| OFClat_R | 3.2841 | 0.0005 |
| Insula_L | 4.5075 | 0.0001 |
| Insula_R | 4.4756 | 0.0001 |
| Cingulate_Mid_L | 3.6854 | 0.0003 |
| Cingulate_Mid_R | 3.8998 | 0.0002 |
| Hippocampus_L | 4.0146 | 0.0001 |
| Hippocampus_R | 3.7427 | 0.0002 |
| ParaHippocampal_L | 3.6645 | 0.0003 |
| ParaHippocampal_R | 3.7434 | 0.0002 |
| Amygdala_L | 4.0651 | 0.0001 |
| Amygdala_R | 3.6975 | 0.0002 |

| | | |
|---|---|---|
| Fusiform_L | 3.5528 | 0.0003 |
| Postcentral_L | 3.5753 | 0.0003 |
| Postcentral_R | 3.8108 | 0.0003 |
| Parietal_Inf_L | 3.3066 | 0.0005 |
| SupraMarginal_L | 3.4942 | 0.0004 |
| SupraMarginal_R | 3.8776 | 0.0002 |
| Paracentral_Lobule_L | 3.4346 | 0.0004 |
| Caudate_L | 3.5091 | 0.0003 |
| Caudate_R | 3.4317 | 0.0004 |
| Putamen_L | 3.6246 | 0.0003 |
| Putamen_R | 3.7300 | 0.0003 |
| Pallidum_L | 3.2837 | 0.0006 |
| Temporal_Sup_L | 3.5406 | 0.0003 |
| Temporal_Pole_Sup_L | 3.7996 | 0.0002 |
| Temporal_Pole_Sup_R | 3.7752 | 0.0002 |
| Temporal_Pole_Mid_L | 3.4469 | 0.0004 |
| ACC_sub_L | 3.6190 | 0.0003 |
| ACC_sub_R | 3.6659 | 0.0002 |
| ACC_pre_L | 4.2652 | 0.0001 |
| ACC_pre_R | 4.4827 | 0.0001 |
| ACC_sup_L | 3.6695 | 0.0003 |
| ACC_sup_R | 3.9976 | 0.0002 |
| N_Acc_L | 3.5404 | 0.0003 |
| N_Acc_R | 3.5088 | 0.0003 |

Each ROI label is in accordance with the automated anatomical atlas 3 (6). L, left hemisphere. R, right hemisphere.

**Table S5.** Brain regions that significantly increased for rapid eye movements during REM sleep.

| ROIs | *Z* values | *P* values |
|---|---|---|
| Supp_Motor_Area_L | 3.2030 | 0.0007 |
| Hippocampus_L | 3.4546 | 0.0004 |
| Hippocampus_R | 3.7187 | 0.0002 |
| ParaHippocampal_L | 3.3809 | 0.0004 |
| ParaHippocampal_R | 3.9307 | 0.0002 |
| Calcarine_L | 3.3290 | 0.0005 |
| Calcarine_R | 3.4544 | 0.0004 |
| Lingual_L | 3.3523 | 0.0005 |
| Lingual_R | 3.4621 | 0.0004 |
| Occipital_Sup_L | 3.2501 | 0.0006 |
| Cerebellum_3_L | 3.2661 | 0.0006 |
| Cerebellum_3_R | 3.1315 | 0.0009 |
| Cerebellum_4_5_L | 3.6211 | 0.0002 |
| Cerebellum_4_5_R | 3.3396 | 0.0005 |
| Cerebellum_6_L | 3.2634 | 0.0006 |
| Cerebellum_6_R | 3.3340 | 0.0005 |
| Cerebellum_9_L | 3.1488 | 0.0008 |
| Vermis_1_2 | 3.2139 | 0.0007 |
| Vermis_3 | 3.5091 | 0.0004 |
| Vermis_4_5 | 3.4872 | 0.0004 |
| Vermis_6 | 3.3777 | 0.0005 |
| Vermis_10 | 3.1045 | 0.0010 |
| Thal_VA_L | 3.3155 | 0.0005 |
| Thal_VA_R | 3.2629 | 0.0006 |
| Thal_VL_L | 3.6201 | 0.0003 |
| Thal_VL_R | 3.6587 | 0.0003 |
| Thal_VPL_L | 3.5085 | 0.0004 |
| Thal_VPL_R | 3.5499 | 0.0003 |
| Thal_IL_L | 3.9072 | 0.0002 |
| Thal_IL_R | 4.2275 | 0.0000 |
| Thal_MDm_L | 3.8869 | 0.0002 |
| Thal_MDm_R | 3.7111 | 0.0003 |
| Thal_MDl_L | 4.0575 | 0.0001 |
| Thal_MDl_R | 3.7768 | 0.0002 |
| Thal_LGN_L | 3.3645 | 0.0004 |
| Thal_LGN_R | 3.2806 | 0.0006 |
| Thal_MGN_L | 3.3011 | 0.0005 |
| Thal_MGN_R | 3.5171 | 0.0003 |
| Thal_PuI_L | 3.2652 | 0.0006 |
| Thal_PuI_R | 3.2704 | 0.0006 |
| Thal_PuM_L | 3.4233 | 0.0004 |
| Thal_PuM_R | 3.5812 | 0.0003 |

| | | |
|---|---|---|
| Thal_PuA_L | 3.6875 | 0.0002 |
| Thal_PuA_R | 3.7081 | 0.0002 |
| Thal_PuL_L | 3.2427 | 0.0006 |
| Thal_PuL_R | 3.3158 | 0.0005 |
| VTA_R | 3.1182 | 0.0009 |
| SN_pc_L | 3.1993 | 0.0007 |
| SN_pc_R | 3.4885 | 0.0004 |
| SN_pr_L | 3.2564 | 0.0006 |
| SN_pr_R | 3.4536 | 0.0004 |
| Red_N_L | 3.4694 | 0.0003 |
| Red_N_R | 3.6793 | 0.0002 |
| Raphe_D | 3.4396 | 0.0004 |

Each ROI label is in accordance with the automated anatomical atlas 3 (6). L, left hemisphere. R, right hemisphere.

**Table S6.** Brain regions that significantly increased for arousals.

| ROIs | *Z* values | *P* values |
|---|---|---|
| Frontal_Sup_2_R | 3.6E+00 | 3.4E-04 |
| Frontal_Mid_2_L | 3.7E+00 | 3.5E-04 |
| Frontal_Mid_2_R | 3.6E+00 | 2.7E-04 |
| Frontal_Inf_Oper_L | 3.7E+00 | 2.4E-04 |
| Frontal_Inf_Oper_R | 3.6E+00 | 1.9E-04 |
| Frontal_Inf_Orb_2_L | 4.3E+00 | 8.3E-05 |
| Frontal_Inf_Orb_2_R | 4.3E+00 | 1.5E-04 |
| Rolandic_Oper_L | 3.2E+00 | 6.8E-04 |
| Rolandic_Oper_R | 3.3E+00 | 5.0E-04 |
| Supp_Motor_Area_L | 3.8E+00 | 2.2E-04 |
| Supp_Motor_Area_R | 3.6E+00 | 2.8E-04 |
| Olfactory_L | 3.9E+00 | 1.9E-04 |
| Olfactory_R | 4.0E+00 | 1.7E-04 |
| Frontal_Sup_Medial_L | 3.5E+00 | 3.4E-04 |
| Frontal_Med_Orb_R | 3.8E+00 | 6.8E-05 |
| Rectus_L | 3.9E+00 | 1.5E-04 |
| Rectus_R | 4.0E+00 | 1.8E-04 |
| OFCmed_L | 3.4E+00 | 3.9E-04 |
| OFCmed_R | 3.5E+00 | 3.0E-04 |
| OFCant_L | 3.9E+00 | 1.5E-04 |
| OFCant_R | 4.0E+00 | 1.6E-04 |
| OFCpost_L | 3.6E+00 | 3.9E-04 |
| OFCpost_R | 3.7E+00 | 9.3E-05 |
| Insula_L | 3.9E+00 | 2.1E-04 |
| Insula_R | 3.7E+00 | 2.5E-04 |
| Cingulate_Mid_L | 3.9E+00 | 2.0E-04 |
| Cingulate_Mid_R | 3.9E+00 | 2.0E-04 |
| Cingulate_Post_L | 4.3E+00 | 9.7E-05 |
| Cingulate_Post_R | 4.0E+00 | 1.4E-04 |
| Hippocampus_L | 4.1E+00 | 1.4E-04 |
| Hippocampus_R | 4.4E+00 | 1.0E-04 |
| ParaHippocampal_L | 4.0E+00 | 1.9E-04 |
| ParaHippocampal_R | 4.3E+00 | 1.2E-04 |
| Amygdala_L | 4.2E+00 | 1.1E-04 |
| Amygdala_R | 5.0E+00 | 3.1E-05 |
| Calcarine_R | 3.7E+00 | 2.0E-04 |
| Cuneus_R | 3.8E+00 | 1.7E-04 |
| Lingual_L | 3.7E+00 | 2.7E-04 |
| Lingual_R | 4.1E+00 | 1.3E-04 |
| Occipital_Sup_L | 3.6E+00 | 2.5E-04 |
| Occipital_Sup_R | 3.2E+00 | 7.4E-04 |
| Occipital_Inf_L | 3.7E+00 | 2.1E-04 |

| | | |
|---|---|---|
| Fusiform_L | 3.8E+00 | 2.2E-04 |
| Fusiform_R | 4.3E+00 | 1.2E-04 |
| Parietal_Sup_L | 3.5E+00 | 3.7E-04 |
| Parietal_Inf_L | 3.4E+00 | 2.8E-04 |
| SupraMarginal_R | 3.1E+00 | 9.2E-04 |
| Angular_L | 3.3E+00 | 5.1E-04 |
| Precuneus_L | 3.6E+00 | 3.1E-04 |
| Precuneus_R | 3.7E+00 | 2.7E-04 |
| Paracentral_Lobule_L | 3.8E+00 | 2.5E-04 |
| Paracentral_Lobule_R | 3.7E+00 | 3.0E-04 |
| Caudate_L | 4.6E+00 | 8.3E-05 |
| Caudate_R | 5.1E+00 | 2.1E-05 |
| Putamen_L | 4.9E+00 | 3.9E-05 |
| Putamen_R | 4.9E+00 | 4.3E-05 |
| Pallidum_L | 4.7E+00 | 7.7E-05 |
| Pallidum_R | 4.6E+00 | 3.0E-05 |
| Heschl_R | 3.3E+00 | 5.8E-04 |
| Temporal_Sup_L | 3.4E+00 | 4.4E-04 |
| Temporal_Sup_R | 3.6E+00 | 3.7E-04 |
| Temporal_Pole_Sup_L | 3.3E+00 | 5.8E-04 |
| Temporal_Pole_Sup_R | 3.5E+00 | 3.3E-04 |
| Temporal_Mid_L | 3.5E+00 | 4.4E-04 |
| Temporal_Mid_R | 3.4E+00 | 3.5E-04 |
| Temporal_Inf_L | 3.9E+00 | 2.4E-04 |
| Temporal_Inf_R | 3.8E+00 | 2.4E-04 |
| Cerebellum_Crus1_L | 4.7E+00 | 6.1E-05 |
| Cerebellum_Crus1_R | 4.3E+00 | 1.2E-04 |
| Cerebellum_Crus2_L | 4.0E+00 | 1.8E-04 |
| Cerebellum_Crus2_R | 4.1E+00 | 1.4E-04 |
| Cerebellum_3_L | 4.6E+00 | 6.3E-05 |
| Cerebellum_3_R | 4.4E+00 | 9.7E-05 |
| Cerebellum_4_5_L | 4.7E+00 | 6.0E-05 |
| Cerebellum_4_5_R | 4.8E+00 | 5.6E-05 |
| Cerebellum_6_L | 4.7E+00 | 6.7E-05 |
| Cerebellum_6_R | 4.7E+00 | 6.1E-05 |
| Cerebellum_7b_L | 3.6E+00 | 2.8E-04 |
| Cerebellum_8_L | 3.7E+00 | 2.6E-04 |
| Cerebellum_8_R | 3.7E+00 | 2.9E-04 |
| Cerebellum_9_L | 3.7E+00 | 2.4E-04 |
| Cerebellum_9_R | 3.5E+00 | 3.6E-04 |
| Cerebellum_10_L | 3.6E+00 | 3.6E-04 |
| Cerebellum_10_R | 3.7E+00 | 2.0E-04 |
| Vermis_1_2 | 4.2E+00 | 8.7E-05 |
| Vermis_3 | 4.0E+00 | 1.4E-04 |
| Vermis_4_5 | 5.3E+00 | 1.7E-05 |

| | | |
|---|---|---|
| Vermis_6 | 5.6E+00 | 8.2E-06 |
| Vermis_7 | 4.8E+00 | 1.2E-05 |
| Vermis_8 | 4.7E+00 | 7.9E-05 |
| Vermis_9 | 4.1E+00 | 1.2E-04 |
| Vermis_10 | 3.5E+00 | 3.5E-04 |
| Thal_AV_L | 5.6E+00 | 7.0E-06 |
| Thal_AV_R | 5.1E+00 | 5.5E-05 |
| Thal_LP_L | 3.8E+00 | 2.0E-04 |
| Thal_LP_R | 3.7E+00 | 2.4E-04 |
| Thal_VA_L | 5.6E+00 | 2.8E-07 |
| Thal_VA_R | 5.9E+00 | 1.4E-06 |
| Thal_VL_L | 5.3E+00 | 7.3E-06 |
| Thal_VL_R | 5.6E+00 | 1.0E-05 |
| Thal_VPL_L | 4.5E+00 | 7.5E-05 |
| Thal_VPL_R | 4.6E+00 | 7.5E-05 |
| Thal_IL_L | 4.6E+00 | 5.1E-05 |
| Thal_IL_R | 4.7E+00 | 1.8E-05 |
| Thal_Re_L | 5.2E+00 | 6.4E-07 |
| Thal_Re_R | 5.3E+00 | 5.5E-08 |
| Thal_MDm_L | 5.7E+00 | 1.6E-06 |
| Thal_MDm_R | 5.6E+00 | 1.5E-06 |
| Thal_MDl_L | 5.6E+00 | 3.2E-07 |
| Thal_MDl_R | 5.3E+00 | 1.3E-07 |
| Thal_LGN_L | 3.2E+00 | 6.3E-04 |
| Thal_LGN_R | 3.9E+00 | 1.3E-04 |
| Thal_MGN_L | 3.3E+00 | 5.1E-04 |
| Thal_MGN_R | 3.6E+00 | 3.1E-04 |
| Thal_PuI_L | 3.5E+00 | 2.3E-04 |
| Thal_PuM_L | 4.0E+00 | 1.5E-04 |
| Thal_PuM_R | 4.4E+00 | 1.1E-04 |
| Thal_PuA_L | 4.4E+00 | 4.6E-05 |
| Thal_PuA_R | 4.2E+00 | 8.6E-05 |
| Thal_PuL_L | 4.1E+00 | 1.0E-04 |
| Thal_PuL_R | 3.8E+00 | 2.1E-04 |
| ACC_sub_L | 3.7E+00 | 3.8E-04 |
| ACC_sub_R | 3.8E+00 | 1.9E-04 |
| ACC_pre_L | 3.5E+00 | 3.4E-04 |
| ACC_pre_R | 3.6E+00 | 2.9E-04 |
| ACC_sup_L | 4.5E+00 | 7.4E-05 |
| ACC_sup_R | 4.1E+00 | 1.4E-04 |
| N_Acc_L | 4.7E+00 | 6.6E-05 |
| N_Acc_R | 5.0E+00 | 2.2E-05 |
| VTA_L | 5.4E+00 | 1.2E-06 |
| VTA_R | 5.0E+00 | 1.9E-06 |
| SN_pc_L | 4.8E+00 | 4.4E-05 |

| | | |
|---|---|---|
| SN_pc_R | 4.9E+00 | 4.9E-06 |
| SN_pr_L | 4.3E+00 | 7.6E-05 |
| SN_pr_R | 4.8E+00 | 3.1E-05 |
| Red_N_L | 4.1E+00 | 1.7E-04 |
| Red_N_R | 3.9E+00 | 1.8E-04 |
| LC_L | 4.2E+00 | 6.4E-05 |
| LC_R | 4.0E+00 | 2.8E-04 |
| Raphe_D | 4.5E+00 | 7.5E-05 |
| Raphe_M | 4.6E+00 | 1.3E-05 |

Each ROI label is in accordance with the automated anatomical atlas 3 (6). L, left hemisphere. R, right hemisphere.

**Table S7.** Brain regions that increased significantly for sleep spindles during light NREM sleep than during slow-wave sleep.

| ROIs | *Z* values | *P* values |
|---|---|---|
| Precentral_L | 3.4296 | 0.0004 |
| Precentral_R | 3.4019 | 0.0004 |
| Frontal_Sup_2_R | 3.1556 | 0.0008 |
| Frontal_Mid_2_R | 3.1447 | 0.0008 |
| Frontal_Inf_Oper_L | 3.4841 | 0.0005 |
| Rolandic_Oper_L | 3.6361 | 0.0003 |
| Rolandic_Oper_R | 3.4432 | 0.0004 |
| Supp_Motor_Area_L | 3.4196 | 0.0004 |
| Supp_Motor_Area_R | 3.3864 | 0.0004 |
| Frontal_Sup_Medial_L | 3.2095 | 0.0007 |
| Insula_L | 3.5361 | 0.0003 |
| Insula_R | 3.6424 | 0.0003 |
| Cingulate_Mid_L | 3.4839 | 0.0003 |
| Cingulate_Mid_R | 3.5164 | 0.0003 |
| Postcentral_L | 3.4082 | 0.0004 |
| Postcentral_R | 3.3369 | 0.0005 |
| SupraMarginal_L | 3.3497 | 0.0005 |
| SupraMarginal_R | 3.3474 | 0.0004 |
| Precuneus_L | 3.2277 | 0.0007 |
| Precuneus_R | 3.2651 | 0.0006 |
| Paracentral_Lobule_L | 3.3886 | 0.0004 |
| Paracentral_Lobule_R | 3.4239 | 0.0004 |
| Putamen_R | 3.5491 | 0.0003 |
| Heschl_L | 3.7458 | 0.0002 |
| Heschl_R | 3.7344 | 0.0002 |
| Temporal_Sup_L | 3.6067 | 0.0003 |
| Temporal_Sup_R | 3.5031 | 0.0003 |
| Temporal_Pole_Sup_L | 3.5521 | 0.0003 |
| Temporal_Pole_Sup_R | 3.3365 | 0.0005 |
| Temporal_Mid_L | 3.4501 | 0.0004 |
| Temporal_Mid_R | 3.1425 | 0.0008 |
| ACC_sup_L | 3.3683 | 0.0004 |
| ACC_sup_R | 3.2987 | 0.0005 |

Each ROI label is in accordance with the automated anatomical atlas 3 (6). L, left hemisphere. R, right hemisphere.

**Table S8.** Brain regions that increased significantly for sleep spindles during slow-wave sleep than during light NREM sleep.

| ROIs | *Z* values | *P* values |
|---|---|---|
| ParaHippocampal_R | 3.2843 | 0.0005 |
| Calcarine_R | 3.7146 | 0.0003 |
| Cuneus_L | 3.3800 | 0.0005 |
| Cuneus_R | 3.5743 | 0.0003 |
| Lingual_R | 3.2666 | 0.0006 |
| Occipital_Sup_L | 3.2750 | 0.0006 |
| Fusiform_R | 3.2916 | 0.0006 |
| Parietal_Sup_L | 3.3117 | 0.0005 |
| Parietal_Sup_R | 3.2652 | 0.0006 |
| Precuneus_L | 3.2913 | 0.0006 |
| Precuneus_R | 3.7949 | 0.0002 |
| Cerebellum_Crus1_R | 3.2078 | 0.0007 |
| Cerebellum_3_R | 3.2160 | 0.0007 |
| Cerebellum_4_5_R | 3.3673 | 0.0005 |
| Cerebellum_6_R | 3.3231 | 0.0005 |

Each ROI label is in accordance with the automated anatomical atlas 3 (6). L, left hemisphere. R, right hemisphere.

**Table S9.** Brain regions that increased significantly for sawtooth waves during REM sleep.

| ROIs | *Z* values | *P* values |
|---|---|---|
| Frontal_Inf_Oper_R | 3.2735 | 0.0005 |
| Calcarine_L | 3.2675 | 0.0006 |
| Lingual_L | 3.2436 | 0.0006 |
| Paracentral_Lobule_R | 3.1408 | 0.0008 |
| Heschl_L | 3.1726 | 0.0008 |
| Vermis_9 | 3.2986 | 0.0005 |
| Thal_VPL_L | 3.1857 | 0.0007 |
| Thal_PuM_L | 3.2213 | 0.0007 |
| Thal_PuL_L | 3.2732 | 0.0005 |

Each ROI label is in accordance with the automated anatomical atlas 3 (6). L, left hemisphere. R, right hemisphere.

**SI References**


1. A. R. Adamantidis, C. Gutierrez Herrera, T. C. Gent, Oscillating circuitries in the sleeping brain. *Nat Rev Neurosci* **20**, 746–762 (2019).
2. D. Picchioni, *et al.*, Autonomic arousals contribute to brain fluid pulsations during sleep. *Neuroimage* **249**, 118888 (2022).
3. S. D. Williams, *et al.*, Neural activity induced by sensory stimulation can drive large-scale cerebrospinal fluid flow during wakefulness in humans. *PLoS Biol* **21**, e3002035 (2023).
4. C. Strik, U. Klose, M. Erb, H. Strik, W. Grodd, Intracranial oscillations of cerebrospinal fluid and blood flows: Analysis with magnetic resonance imaging. *Journal of Magnetic Resonance Imaging* **15**, 251–258 (2002).
5. R. Berry, S. Quan, A. Abreu, Manual for the Scoring of Sleep and Associated Events: Rules, Terminology and Technical Specifications. *American Academy of Sleep Medicine* **Version 2.6** (2020).
6. E. T. Rolls, C.-C. Huang, C.-P. Lin, J. Feng, M. Joliot, Automated anatomical labelling atlas 3. *Neuroimage* **206**, 116189 (2020).
